# The amino acid sequence determines protein abundance through its conformational stability and reduced synthesis cost

**DOI:** 10.1101/2023.10.02.560091

**Authors:** Filip Buric, Sandra Viknander, Xiaozhi Fu, Oliver Lemke, Jan Zrimec, Lukasz Szyrwiel, Michael Mueleder, Markus Ralser, Aleksej Zelezniak

**Affiliations:** - Department of Biology and Biological Engineering, Chalmers University of Technology, Kemivägen 10, SE-412 96, Gothenburg, Sweden; - Department of Biochemistry, Charité – Universitätsmedizin Berlin, 10117 Berlin, Germany; - Department of Biotechnology and Systems Biology, National Institute of Biology, Večna pot 111, SI1000 Ljubljana, Slovenia; - Core Facility High Throughput Mass Spectrometry, Charité – Universitätsmedizin Berlin, 10117 Berlin, Germany; - Institute of Biotechnology, Life Sciences Centre, Vilnius University, Sauletekio al. 7, LT10257 Vilnius, Lithuania; - Randall Centre for Cell & Molecular Biophysics, King’s College London, New Hunt’s House, Guy’s Campus, SE1 1UL London, UK

**Author notes:** These authors contributed equally (Filip Buric, Sandra Viknander).

**Keywords:** proteome, protein sequence, protein expression, protein stability, deep learning, language models, explainable machine learning, molecular dynamics

## Abstract

Understanding what drives protein abundance is essential to biology, medicine, and biotechnology. Driven by evolutionary selection, the amino acid sequence is tailored to meet the required abundance of proteomes, underscoring the intricate relationship between sequence and functional demand. Yet, the specific role of amino acid sequences in determining proteome abundance remains elusive. Here, we demonstrate that the amino acid sequence predicts abundance by shaping a protein’s conformational stability. We show that increasing the abundance provides metabolic cost benefits, underscoring the evolutionary advantage of maintaining a highly abundant and stable proteome. Specifically, using a deep learning model (BERT), we predict 56% of protein abundance variation in *Saccharomyces cerevisiae* solely based on amino acid sequence. The model reveals latent factors linking sequence features to protein stability. To probe these relationships, we introduce MGEM (Mutation Guided by an Embedded Manifold), a methodology for guiding protein abundance through sequence modifications. We find that mutations increasing abundance significantly alter protein polarity and hydrophobicity, underscoring a connection between protein stability and abundance. Through molecular dynamics simulations and *in vivo* experiments in yeast, we confirm that abundance-enhancing mutations result in longer-lasting and more stable protein expression. Importantly, these sequence changes also reduce metabolic costs of protein synthesis, elucidating the evolutionary advantage of cost-effective, high-abundance, stable proteomes. Our findings support the role of amino acid sequence as a pivotal determinant of protein abundance and stability, revealing an evolutionary optimization for metabolic efficiency.

## Introduction

The intricate interplay between protein synthesis and degradation defines intracellular protein levels, with implications for therapeutic strategies, as well as efficient protein and cellular engineering. The complex regulation of protein homeostasis suggests that multiple factors contribute to the overall proteome makeup, with the evolutionarily encoded sequence potentially playing a pivotal role in proteome composition. For instance, protein synthesis is strongly regulated at the initiation step ^1,2^, whose rate varies broadly between mRNAs, depending not only on the transcript sequence features but also on the amino acids at the N-terminal ^3,4^. In bacteria, the amino acid composition of the C-terminal is a strong determinant of protein degradation rates, explaining a wide range of protein abundances ^5,6^. These, along with the multiple mechanisms of post-translational regulation ^7,8^, suggest that this rather tight regulation occurs at the degradation level and is encoded, at least partially, in the amino acid sequence. Empirically, amino acid composition and sequence features were seen to correlate with protein abundance ^9–11^, transcending mere codon composition influences on protein abundance^12^. While the importance of protein sequence in determining abundance is recognised, the quantitative relationship between sequence and abundance remains elusive, as does the link between the evolutionary mechanisms that underlie this relationship.

On a broader scale, proteins situated as central players in cellular processes or as critical nodes in interaction networks often exhibit higher abundances ^13^. Evolutionarily, these highly abundant proteins face stringent constraints, evolving at a slower pace due to their potential large-scale impact on cellular fitness ^14,15^. Remarkably, the conservation of steady-state protein abundances spans across diverse evolutionary lineages, ranging from bacteria to human ^16–18^. Theoretical models suggest that increasing protein abundance slows evolution due to reduced fitness, with the least stable proteins adapting the fastest ^19^. Yet, under strong selection, proteins can evolve faster by adopting mutations that enhance stability and folding ^20^. Experimental evidence also suggests that a protein’s capacity to evolve is enhanced by the mutational robustness conferred by extra stability ^21–23^, meaning that protein stability increases evolvability by allowing a protein to accept a broader range of beneficial mutations while still folding to its native structure. Thermostability gains of highly expressed orthologs are often accompanied by a more negative ΔG of folding, indicating that highly expressed proteins are often more thermostable ^24^, as often explained by the so-called misfolding avoidance hypothesis (MAH), because stable proteins are evolutionarily designed to tolerate translational errors ^25–27^. On the contrary, several empirical studies revealed no substantial correlation between protein stability and protein abundance ^28,29^. Likewise, the overall cost (per protein) of translation-induced misfolding is low compared to the metabolic cost of synthesis ^30,31^, suggesting that MAH does not explain why highly abundant proteins evolve slower ^29^. On the other hand, cells may have fine-tuned protein sequences to balance their functional importance with the metabolic costs they incur, reflecting an optimisation between functional necessity and energy efficiency ^32–34^. Given the intricate interplay of evolutionary constraints, protein stability, abundance, and metabolic cost, it still remains unclear how cells evolved their sequences to strike an optimal balance between functional demands of proteome and cellular fitness associated with synthesis and maintenance of protein abundance.

In this study, we explored the relationship between a protein’s amino acid sequence and its abundance. Using a deep neural network transformer (BERT) trained on data from 21 proteome studies, we could predict over half of the protein copy number variation (R^2^ _test_ = 56%) in *Saccharomyces cerevisiae* based solely on amino acid sequences. Delving into the neural network’s self-attention mechanism to understand which protein sequence features are predictive of their abundances, we revealed that the network indirectly identified specific physicochemical properties inherently encoded in amino acid sequences related to a protein’s conformational stability. We then introduced MGEM (Mutation Guided by an Embedded Manifold) to probe sequence space and found that abundance-enhancing mutations notably affected protein polarity and hydrophobicity, hinting at a stability-abundance connection. Molecular dynamics simulations further confirmed the enhanced stability of abundance-increasing mutants. Using a proteomics experiment in yeast, we revealed that mutant protein remained more abundant over the course of yeast growth phases compared to a wild type variant. Importantly, we found that mutants with increased abundance had lower amino acid synthesis costs than their native versions, underscoring the fitness benefits of abundant, stable proteins. Our research shows that the amino acid sequence is a key factor influencing intracellular protein levels. This is achieved by boosting protein stability, which is driven by cost-effective amino acid substitutions, providing evolutionary benefits by reducing the metabolic costs of protein synthesis.

## Results

### The amino acid sequence is predictive of protein abundance

To investigate the relationship between amino acid sequence and protein abundance, we used a compendium of 21 experimental systematic quantitative studies employing mass spectrometry and microscopy to estimate absolute protein abundances of over 5000 proteins (copy numbers per cell) in *Saccharomyces cerevisiae* grown predominantly in the exponential phases across multiple conditions essentially capturing proteome variation ^35^. The gene-wise dynamic range of protein abundances spanned an average of 5 orders of magnitude, while individual protein expression values for 95% of proteins varied within only one relative standard deviation (RSD) across all experimental conditions (Figure S1). A similar phenomenon has been observed previously with mRNA levels encoded in the DNA sequences ^36,37^. This result suggests that individual protein expression across experimental conditions primarily fluctuates around a specific expression value, suggesting its deterministic nature.

Next, to investigate the relationship between amino acids and intracellular protein levels, we formulated a regression problem by utilising protein sequences to model protein abundance values. To learn sequences, we chose the Bidirectional Encoder Representations from Transformers (BERT) architecture ^38,39^, which allows for transparency in weighing the contributions of amino acid residues on protein levels and provides insights into the most relevant sequence features the model uses ^39–41^ to make predictions about protein abundances, using an intrinsic attention mechanism^42^. Due to deep learning’s need for extensive training data and the yeast dataset’s limited size, we used repeated measurements (up to 21 sequence copies from all experiments in the dataset) to account for inter-experimental variability (equivalent to regression with replicates). Our augmented dataset included 199,206 training examples, with 10% of random sequences uniquely chosen for validation during model training and 10% for a hold-out test during final model evaluation (Methods M1). By training BERT from scratch, we found that the model predicts 56% of protein abundance variation (R^2^ = 56% on a holdout test set) using only an amino acid sequence as input, suggesting that the sequence predominantly encodes protein abundance. In contrast, the model predictions failed completely when performing a randomization test with shuffled sequences (R^2^ = −73%, Figure 1A inset), confirming that the model relies on residue interdependencies in a sequence rather than simply learning amino acid frequencies when predicting protein levels. Further analysis confirmed that amino acid frequency is uniformly distributed across the entire dynamic range of protein abundances, with a mean CV of 7% over abundance deciles (Figure S1D), supporting the neural network’s ability to pick up information encoded in the sequence.

**Figure 1.**
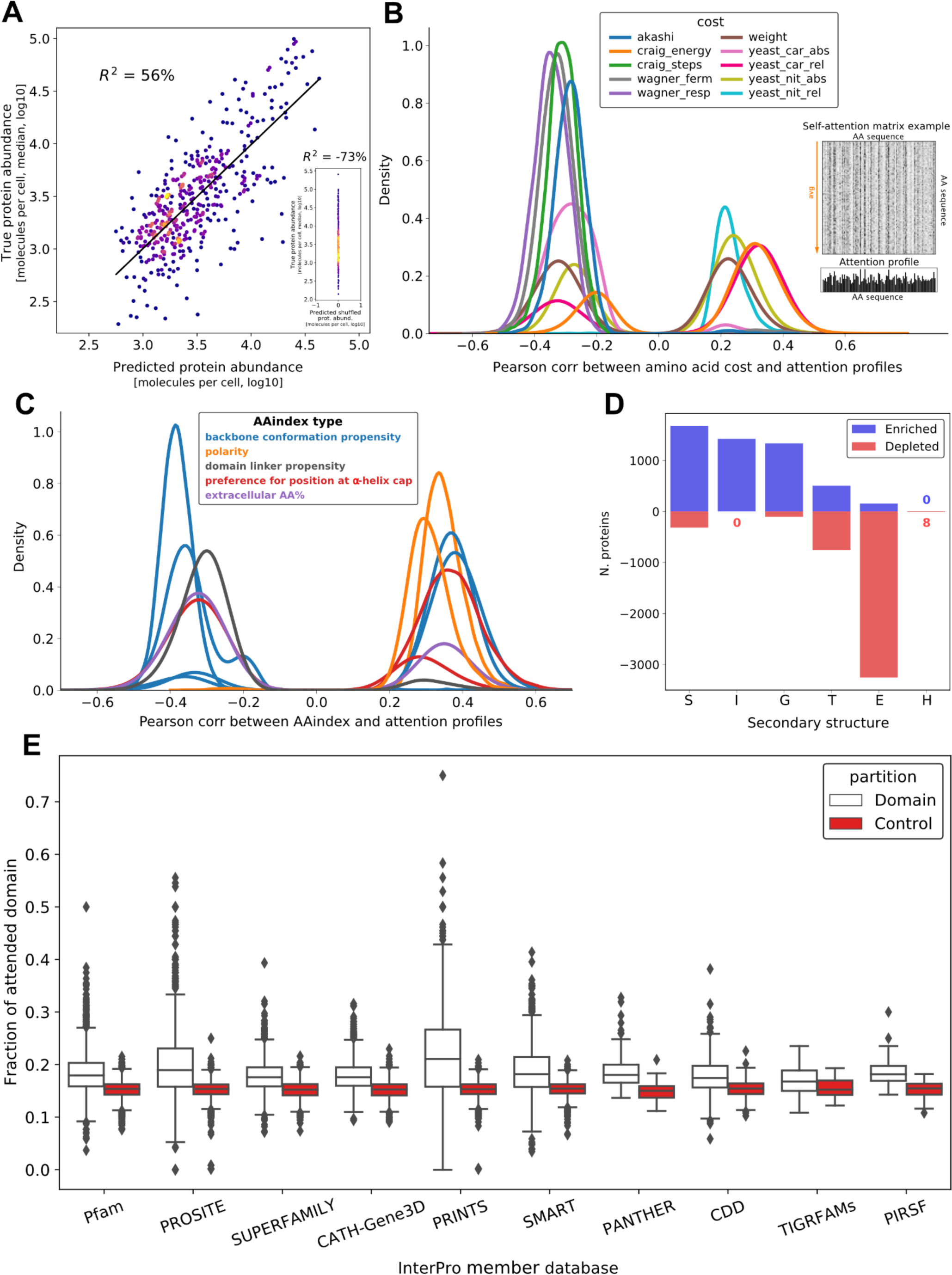
The amino acid sequence is predictive of protein abundance. **A)** BERT performance on a hold-out test set, coloured by density. **Inset:** Random prediction control using shuffled versions of the test sequences. The poor performance on randomized input, effectively predicting a single value, demonstrates that the model has learned sequence structure and not amino acid frequencies. **B)** Attention profiles correlate with amino acid metabolic costs (see also Table S1 for full description). Shown are distributions across all sequences of maximum (absolute) Pearson correlations of any attention profile with p-value < 1e-5. **Inset**: A BERT attention matrix example (top) and derived attention profile (bottom) for a short sequence. Attention matrices consist of directional association weights between pairs of residues, normalized as a percentage. The profiles were obtained by averaging along the “attends-to” axis, as the “attended-by” variation is generally more informative, resulting in one-dimensional attention profiles. **C)** Attention profiles correlate with 10 non-redundant AAindex variables (colored by index type), showing that profiles capture information pertaining to backbone conformation, physicochemical properties, domain linkage, and secondary structure. While some AAindex types correlate with attention profiles both positively and negatively (e.g. backbone conformation), individual AAindex variables within these types are overall either positively or negatively correlated. The categories shown span AAindex variables that are both positively and negatively correlated with attention. **D)** Proteins are split into two subpopulations of sequences with high attention values (z-score > 1) that are either enriched in turns and helices (S, I, G, and T in DSSP notation) and, to a lesser extent, extended strand (E), or largely depleted in extended strand (E) and turn (T), as assessed with one-sided hypergeometric tests (p-value < 0.05). **E)** Overlap of attention patterns with protein domains from the yeast InterPro database, grouped by member databases. The attention coverage of domains (fraction overlapping with attention profiles) is significantly higher than control for 10 out of 12 member databases (Wilcoxon two-sided signed-rank test, p-value < 0.05), with the highest coverage in PRINTS and PROSITE.

### The attention mechanism identifies sequence and structural features linked to protein abundance

Next, we wanted to interpret the features learned by the transformer which explain protein abundance. Models generated by deep neural networks are often difficult to interpret ^43^, however the self-attention mechanism used by transformers has been shown to match multiple physicochemical properties and substitution likelihoods of amino acids ^40^. To increase interpretability of the model as a map of sequence-to-protein abundances, we trained the model from scratch, as opposed to fine-tuning pretrained large protein language models ^44–47^. Protein language embeddings, including sequence representations learned from structural models ^48^, have been shown to have limited generalization to all protein functions and properties ^49,50^, thus making it difficult to use for generalized interpretation. Instead, by training the model from scratch in a regression setting, we ensured that our model learned relevant sequence representations related to protein abundance, easing interpretation. Thus, we next attempted to identify abundance-related links to physicochemical protein features using the attention values derived from yeast protein sequences. We extracted the attention weights of each input sequence and obtained one-dimensional per-residue attention profiles, which reflected the average percentage of attention that each residue receives from all others in the sequence when making the corresponding abundance prediction (see Figure S2 and Methods M2).

To examine the determinants of protein abundance, we first correlated attention profiles with amino acid costs ^51^ (Methods M3), as amino acid synthesis cost is known to be a determinant of protein abundance ^32,52–54^. The strongest correlations were found between attention profiles and the energetic cost of amino acids (*craig_energy*) ^55^ averaged over all proteins (mean Pearson’s r = 0.32, BH adj. p-value < 1e-5). Conversely, anticorrelations were observed with synthetic cost under both respiratory and fermentative growth (*wagner_resp*, *wagner_ferm*, respectively) ^54^ as well as the number of synthesis steps (*craig_steps*) ^55^ (mean Pearson’s r = −0.35, −0.33, and −0.31, respectively, BH adj. p-value < 1e-5). Additionally, some of the systemic costs introduced by Barton et al. ^51^ using genome-scale flux balance analysis calculations ^56^ showed positive and negative correlations with attention, such as the impact of the relative change of the amino acid requirement on the minimal intake of glucose (*yeast_car_rel*, mean Pearson’s r = 0.32 over 1855 proteins and −0.33 over 705 proteins) and the absolute change of the amino acid requirement on the minimal intake of ammonium (*yeast_nit_abs*, mean Pearson’s r = 0.25 over 1833 proteins and −0.28 over 1165 proteins, Figure 1B and Table S1). A negative correlation with synthesis cost implies that the model assigns more weight to "cheaply" synthesized amino acids. In contrast, a positive correlation with energy cost implies paying attention to more energy-rich amino acids when predicting protein abundance. We stress that the correlations reported here do not directly link cost values to the predicted abundance, but rather underline the relevant latent features learned from protein sequence that the model picked up intrinsically prior to mapping sequence to protein levels.

Based on our observation that amino acid frequency is uniformly distributed across the entire dynamic range of protein abundances (Figure S1D), we did not expect to find specific single amino acids that would determine abundances. Instead, we hypothesized that the neural network would capture higher-order interactions important for structural and functional protein features. Thus, we correlated attention profiles with a subset of 18 non-redundant AAindex values representing various physicochemical and biochemical protein properties ^57^ (see Methods M4). We identified significant correlations with measures of backbone *conformation propensity* (both positively and negatively correlated indices, with the strongest mean correlations being 0.38 and −0.38, respectively, p-value < 1e-5), *preference for position at α-helix cap* (both positively and negatively correlated indices, with the strongest mean correlations per sequence being 0.37 and −0.33, respectively, p-value < 1e-5), *polarity* (highest mean correlation = 0.35, p-value < 1e-5), *domain linker propensity* (mean correlation = −0.31, p-value < 1e-5), and *the composition of extracellular domains seen in membrane proteins* (two protein subpopulations, one with mean correlation = 0.36, the other with mean anticorrelation = −0.33, p-value < 1e-5) (Figure 1C, see Tables S2 and S3 for a detailed description). Physicochemical properties of amino acids, such as polarity, have been shown to affect translation speed ^11^ and protein stability ^58^. The correlations with backbone conformation and preference for α-helix cap indicators suggest a link to secondary structure, while the correlation with domain linker propensity points to the model having learned to some extent the boundaries of domain separation.

We next assessed the connection between secondary structure and attention profiles by analyzing the enrichment of per-residue DSSP annotations ^59,60^ in high-attention positions using AlphaFold2 - generated^48^ structures for 4745 yeast proteins. We counted the annotations at positions with attention profile z-scores > 1 and compared them to background annotation counts across all proteins (using one-sided hypergeometric tests for enrichment and depletion, p-value < 0.05) (Methods M5). The results showed that attention values were enriched in turns and helices (S, I, G, and T in DSSP notation) but depleted in extended strands (E) for most proteins (3254 proteins) (Figure 1D). For turns (T), the protein subpopulations were more evenly split, with this structure enriched in 505 proteins and depleted in 754 proteins. These findings suggest that helical structures may be implicated in protein abundance, while the contribution of turns and sheets towards the model prediction may be more complex.

As structural properties imply function, we also investigated whether abundance-driven attention specifically focuses on any functional regions of protein sequences. We examined the extent to which the attention patterns cover the domains from the *S. cerevisiae* InterPro ^61^ database. To allow for comparison with controls, we focused only on domains with a length less than half of the protein sequence, analyzing a total of 18,000 domains (Methods M6). For 10 out of 12 member databases, domains were significantly more covered by high attention than random regions of the same length (Wilcoxon two-sided signed-rank test, adj. p-value < 0.05) (Figure 1E). The results are particularly striking as our BERT model was trained from scratch, not pre-trained on domains as in the study by Rao et al. ^39^. We next performed a GO enrichment analysis on proteins with well-covered domains (chosen as at least 30% domain length overlapping with attention patterns, well above the random control), a total of 832 domains in 517 proteins (Methods M7). From the enriched terms, GO-slim terms were produced for summarization (Table S4). The enriched (Hypergeometric test, adj. p-value < 0.05) biological processes are diverse and, among others, include translation, protein folding, modification, and metabolic processes; the molecular functions include cytoskeletal protein binding, unfolded protein binding, DNA and RNA binding, transmembrane transporter activity and others. This variety points at widespread domain patterns to which the model attends across different protein classes rather than specific functional motifs, which hints at the role of sequence across the entire proteome. On the technical side of the attention mechanism itself, it is interesting to note that domains were predominantly captured by a single (and deeper) network layer (Figure S3).

### Navigating the sequence space to control protein abundance

We next hypothesized that our model could facilitate precise control over protein abundance by introducing targeted changes to the protein sequence. To achieve this, we developed a Mutation procedure Guided by an Embedded Manifold (MGEM), which enables us to navigate the BERT model’s embedded sequence manifold and perform individual amino acid substitutions that increase abundance. The approach involves traversing a uni-dimensional UMAP projection of the BERT encoder’s high-dimensional embedded space, which assigns a scalar importance value to each residue in a sequence based on its impact on protein abundance (i.e. as determined by both position and amino acid that the model learned) (Figure 2A). MGEM substitutes low-importance residues in a starting wild type sequence with high-importance residues from a set of guide sequences selected based on their topmost abundance levels (Figure 2B, see details in Methods M8 and M9). Thus, by borrowing important amino acids (as measured by their order in the UMAP projection) from highly abundant proteins, the modified sequence is “moved” towards higher abundance. This is based on the posited property of the high-dimensional BERT embedded space by which the sequence representations are approximately ordered (or “ranked”) according to the target value (Figure 2A). The per-residue importance values obtained with UMAP are a good approximation of this ordering (Spearman’s ρ = 0.8, p-value < 1e-16) (Figure 2C), enabling the sorting of all residues on a univariate scale that spans all sequences, according to their importance towards prediction (see Methods M8). Our novel method relies on the learned relationship between sequences and only minimally changes wild types by deterministically substituting the individual amino acids directly related to the abundance, without relying on probabilistic or stochastic optimization searches.

**Figure 2.**
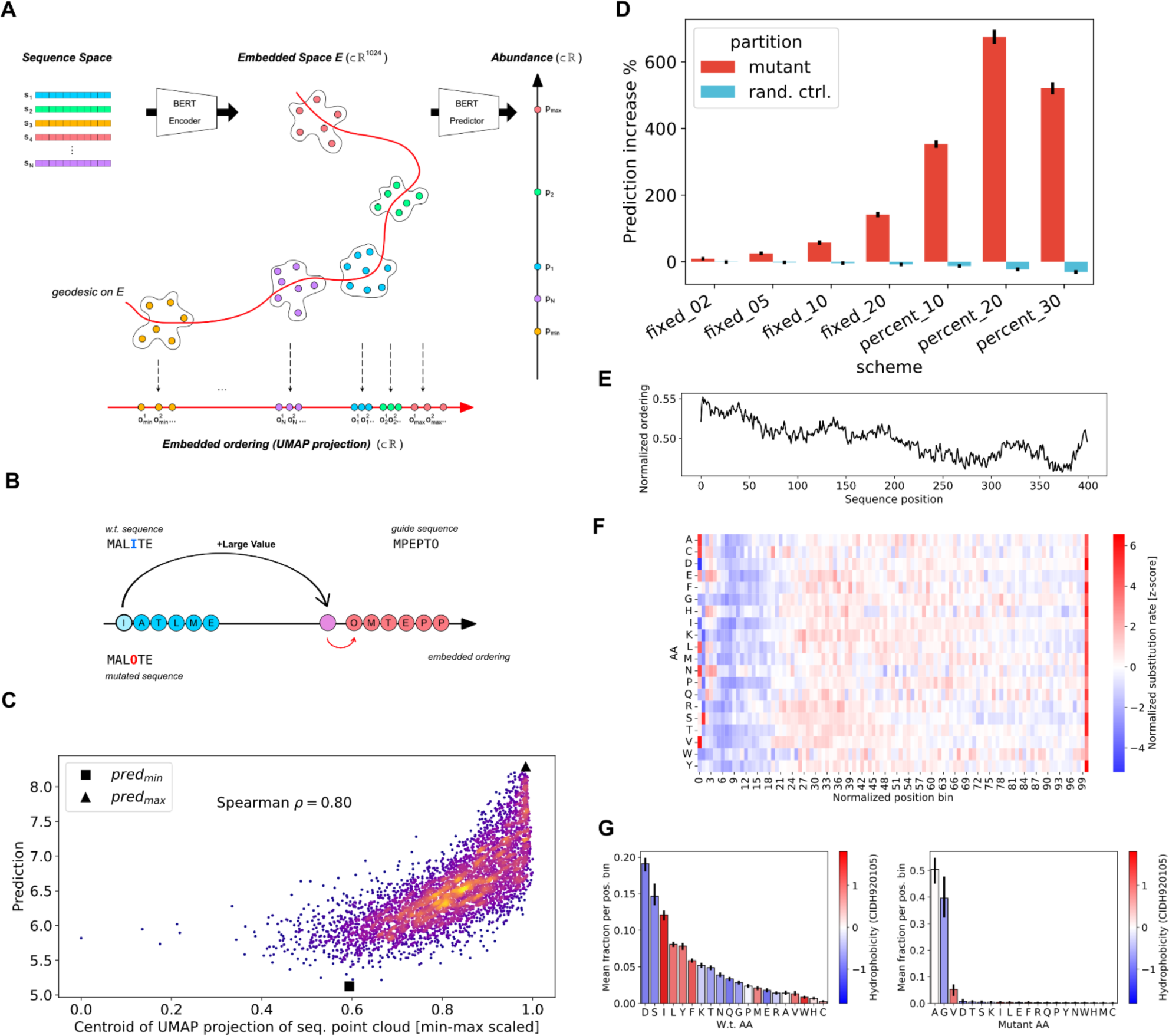
Navigating the sequence space to control protein abundance through guided mutation. **A)** Conceptual illustration showing the posited structure of the BERT encoder embedded space and the embedded ordering construction that supports our guided mutation procedure. The encoder maps each residue in a sequence to a high-dimensional point in the embedded space *E* and sequences thus appear as point clouds. From a point cloud, a thin feedforward predictor yields an abundance prediction. The embedded space is posited to be structured in such a way as to allow a “traversal” of the point clouds, on a path or *geodesic* between all points (curved red line) connecting the points that are part of the lowest abundance sequences to the highest, in an increasing order of predicted values. This path in high-dimensional space is approximated with a parametric UMAP projection from the embedded space *E* to a single dimension, thus giving a simple linear ranking (or ordering) *o_i_^j^* for each residue *j*, in each sequence *i*. This ranking serves to indicate the global weight of a given residue towards the final prediction, compared with all other residues across all sequences. **B)** Simplified illustration of MGEM (mutation guided by embedded manifold) procedure, which takes advantage of the global embedded order value (“importance”) obtained for each residue, across all sequences. The residues with the lowest order value in a sequence are selected for substitution (the “I” residue at position 4 in the illustration) and their order values are increased by a large amount, as a higher value would yield a greater abundance. As we do not have an inverse mapping from this new value to an amino acid, we find the substitute by taking “inspiration” from guide sequences, chosen as the top 10 highest abundance sequences. The residue with closest ordering value to the newly increased value (“O” in the example) is taken and this amino acid replaces the original one in the wild type sequence. **C)** The UMAP projection is a good approximation of the embedded manifold, as it generally correlates well with abundance (Spearman p-value < 1e-308) (the plot is colored by density). Each point corresponds to the centroid of a sequence point cloud, projected through the learned UMAP function. The horizontal axis is normalized to the smallest and largest values in the set of projected points. The centroid of the lowest abundance sequence is marked with a black square and that of the highest abundance sequence with a black triangle. The approximation is worse for lower abundance sequences, as the red square should have appeared as the minimum ordering value. **D)** Predicted abundance increase on sequences mutated with MGEM (black bars showing averages, with 95% confidence intervals). An increasingly higher number of residues with lowest ordering (2, 5, 10, 20 residues, as well as 10%, 20%, and 30% of the sequence) were selected in each scheme shown in the figure. The highest overall increase occurred for the scheme consisting of mutating the 20% lowest-order residues. All schemes showed significantly higher values than random control (blue), which on average decreases predicted abundance. **E)** The most important part of the sequence for the model is the N-terminus, as measured by the embedded ordering value, here normalized to the inverse ranking of residue values (as the relative order is the important information) divided by sequence length. The plot shows the average such profile for sequences of length 200 to 400, the profiles of which were upsampled by linear interpolation to maximum length. **F)** The high importance of the N-terminus for abundance leads to fewer residues being mutated by MGEM, as a consequence of the embedded ordering values (shown in F). Except for the first few positions in the sequence, most amino acids in the leading 20% of the sequence are generally untouched (the leading M is avoided by MGEM). The plot shows for each amino acid the normalized MGEM substitution rate over sequence length bins spanning the leading 30% of sequences (computed over all sequences and mutation schemes). The position has been normalized to sequence length and binned to 2 decimals (resulting in 100 bins). For each amino acid, the number of times MGEM has replaced it in a bin was divided by the wild type count of that amino acid in the same bin. The z-scores of these values were obtained separately for each amino acid. **G)** Average fraction of wild type (left) and MGEM mutant (right) amino acid over the leading 30% of all mutated sequences (error bars showing 95% confidence intervals). The amino acids are colored by their normalized hydrophobicity ^62^, which highlights the overall mutation shift toward more hydrophobic proteins. The binning was performed as in F), i.e. over 30 of the position 100 bins for each sequence.

We next performed a series of *in silico* sequence perturbation experiments by introducing substitutions that would increase protein abundance. This was done across the entire set of protein sequences, in different substitution schemes, each consisting of changing a given number of lowest importance residues per sequence (a fixed number of 2, 5, 10, and 20 residues, as well as 10%, 20%, and 30% of residues in each sequence). We observed that MGEM enables control of target values (protein abundance) significantly more than a random control (paired t-test, adj. p-value <1e-16 for all schemes) in which a random set of residues of the same size as the MGEM set for the given scheme was selected and mutated to random amino acids (Figure 2D). Indeed, on average, random mutations yielded a decrease in protein abundance. The greatest MGEM increase was obtained when mutating 20% of the sequence, achieving an average 675% predicted abundance increase.

By inspecting MGEM mutants, we discovered that in terms of sequence position, the N-terminus is the most important for abundance prediction. The average wild type embedded ordering (importance) profile peaks over the leading 20% of the sequence (Figure 2E), and as a consequence of the MGEM selection process, results in most amino acids being left unchanged in this region (Figure 2F). Additionally, there is a much shorter hotspot of frequently mutated amino acids at the very last positions of the C-terminus. In accordance with other studies ^3,4^, this would suggest that the N-terminus is generally evolutionarily optimized for expression efficiency. Indeed, the composition of the first 30% of sequences significantly differs from the composition of the full sequences (one-sided hypergeometric test, p-value < 1e-3), with the leading region enriched in Ala (A), His (H), Met (M), Pro (P), Gln (Q), Arg (R), Ser (S), Thr (T) (Table S5). The observation that distributions of substituted amino acids differ from the above (some are replaced uniformly across the entire sequence length) is another indication of the role of both the position and the nature of the amino acid. In terms of replacement amino acids, we observed that the vast majority are A, G, and V (Figure 2G). In terms of physicochemical AAindex variables, mutants show significant perturbations (paired t-test, p-value < 1e-80) (see Table S6 and Figure S4), especially in indices that describe *polarity* (specifically amphiphilicity, with a 19% average decrease), *backbone conformation propensity* (with the largest index average decrease by 18% and the highest average index increase by 9%), and in the *preference for position at α-helix cap* (average decrease by 5%), which suggests a change in the likely secondary structure and a shift towards higher hydrophobicity in the mutants.

### Highly abundant proteins show greater conformational stability at a lower metabolic cost

Mutational analysis from MGEM indicates increased protein abundance primarily from non-polar A, G, V amino acid substitutions (Figure 2G). Alanine is known to stabilize helices while glycine varies in its effects ^63^. Glycine can enhance stability in β-turns ^64^. Valine is common in thermophilic proteins ^58^, and both alanine and valine substitutions often show similar helix impacts ^65^. Cysteine, infrequently substituted by our procedure (Figure 2G), is vital for stability due to its potential for disulfide bridge formation ^66^. Likewise, it has been observed that highly expressed proteins are often more thermostable ^24,67^. Using our method which allows for mutations that increase protein abundance, we sought to determine if the model-learned sequence to abundance mapping is linked to overall protein stability. To corroborate this, we applied molecular dynamics (MD) simulations to 100 pairs (mutant and wild types, WTs) of non-membrane yeast proteins (Figure 2D, 20% mutation regime). Both mutated and their original WT versions were modeled using AlphaFold2 structures (Methods M10) and molecular systems were simulated for 100 ns. While our model does account for entire protein abundance variation (Figure 1A), there is a risk that introduced mutations could destabilize proteins. Therefore, we only considered WT and mutant pairs that converged at the end of the simulation trajectory (Methods M10) considering ∼46% of the simulations in our subsequent analyses. To quantify the degree of protein backbone conformational changes, we started by first comparing the fluctuations of atomic positions, expressed as the standard deviation of residue alpha carbons across the entire course of the MD trajectory (root mean square fluctuations, RMSF) between mutant and WT sequences. 33% of converged systems showed significantly lower RMSF in comparison to WT proteins (Wilcoxon rank sum test, adj. p-value < 1e-2) (Figure 3A, Figure S5). Decreases in protein backbone fluctuations might be a sign of protein stabilization^68–70^. 59% of atomic fluctuations of highly abundant mutants were at least 2 standard deviations lower than the corresponding positions of the WT trajectory (Figure 3B). About 81% of mutations had no direct impact on atomic fluctuations, i.e. we observed changes in fluctuations in residues as high as two standard deviations away from corresponding WT positions with no mutations, suggesting that changes in atomic fluctuations caused by abundance-changing mutations affect overall global protein dynamics, rather than just local residues (Figure 3C).

**Figure 3.**
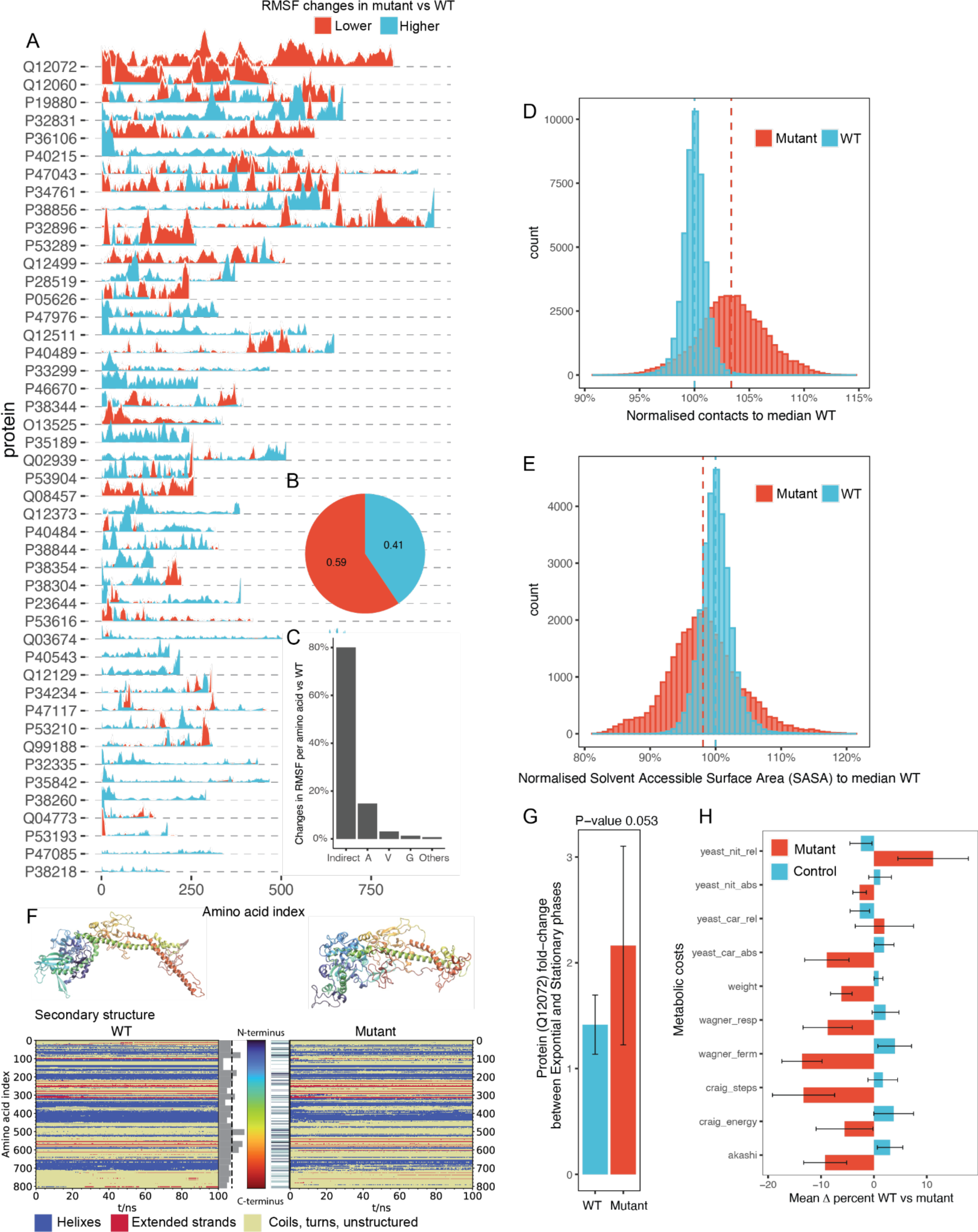
Abundant proteins exhibit higher conformational stability and are synthesized at a lower cost. **A)** Root mean square fluctuations between abundance-increasing mutants and wild type (WT) structures over 100 ns of molecular dynamics trajectory. **B)** Fraction of atomic fluctuation that are at least 2 standard deviations lower in mutant (red) vs wt (blue). **C)** Fraction of total significant (absolute z-score > 2) changes in RMSF per introduced mutation. Indirect denotes the regions of protein sequence with no mutations. **D)** Comparison of contacts between WT and abundance-increasing mutants. Normalization is done with reference to WT using frames after half of the 100 ns trajectory, contacts are considered at 8Å proximity of carbon backbone (Methods M11). **E)** Comparison of solvent accessible solvent ares (SASA) between WT and abundance-increasing mutants. Normalization is done with reference to WT using frames after half of the 100 ns trajectory. **F)** Structure (top) and DSSP plot (bottom) of the wild type (left) and the mutant (right) of IOC2 yeast protein. The structures represent the last frame of the respective simulation (100 ns). The coloring denotes the amino acid index as shown by the colorbar in the center (N-terminus: blue to C-terminus: red). In the DSSP plot, helical structures are highlighted in blue, extended structures in red and everything else (e.g. coil, turn, unstructured) in yellow. The bar plot represents the mutation rate per ∼32 amino acids per bar; the dashed line represents the average mutation rate per bar. On the right hand side the mutated spots are highlighted. **G)** Ratios of IOC2 (UniprotID: Q12072) peptides between exponential and stationary phases in WT and mutant strains. The experiment was performed in biological triplicates (Methods M12). **H)** MGEM reduces protein cost. The average sequence costs of mutants obtained with MGEM (20% mutated sequence) show significant overall decrease compared with random control (paired t-test, p-value < 1e-308), particularly in terms of synthesis costs (see also Table S7). The exceptions were two systemic costs from Barton et al. ^51^, one having the lowest correlation with attention (12% cost increase on average), and the other having both weakly positively and negatively correlated subpopulations (2% cost increase on average).

Although large structural changes from mutations can destabilize proteins ^68,71^, backbone conformational changes do not directly indicate protein stability. To delve deeper, we examined intermolecular interactions, specifically the number of contacts between neighboring amino acids (Methods M11). Stable proteins with robust hydrophobic cores generally have more native contacts^72^. In our comparison, 84% of the high-abundance mutants exhibited significantly more contacts than their wild types (Wilcoxon rank sum test, adj. p-value < 1e-4) (Figure 3D, Figure S6). Proteins that easily denature expose their hydrophobic core, resulting in lost hydrophobic interactions and increased solvent accessibility^68,73,74^. Investigating the effects of A, G, V substitutions on hydrophobic cores, we computed the Solvent Accessible Surface Area (SASA) for all proteins. We found a significant decrease (Wilcoxon rank sum test, p-value < 1e-4) in SASA for abundance-increasing mutants versus wild types, supporting our hypothesis (Figure 3E).

Next, we closely examined the dynamic effects of mutations on the IOC2 protein (UniprotID: Q12072) based on its top decreased RMSF (Figure 3A). Although the mutant and WT IOC2 started similarly, they diverged dynamically over 100 ns of simulation (Figure 3F, Figure S7). The stable core, largely less mutated, differed from the more mutated C-terminal region (Figure 3F, bar plot). A notable change was the breaking of an alpha-helix in the mutant, enabling the C-terminus to fold closer to the protein core. This change led to an increase (WT: 53.0%, mutant: 59.9%; Mann-Whitney U test, p-value < 1e-16) in the median unstructured secondary structure (Figure 3F, DSSP) but formed a more compact shape than its WT counterpart. Despite imperfect alignment in the C-terminal region, an overall increase in hydrophobicity is seen in the mutant (mean −0.07 with the WT vs. 0.17 with the mutant, Mann-Whitney U test p-value < 1e-4), reflected in a reduced RMSF (Figure 3A, Figure S5). To experimentally validate whether the abundance-increasing mutations could potentially stabilize protein expression *in vivo,* we performed an experiment in *S. cerevisiae* by comparing the changes in protein expression between exponential (E) and stationary (S) phases. Specifically, we genetically replaced the native WT variant with the synthetically mutated IOC2 protein (Methods M12). Using a liquid chromatography-coupled mass spectrometer (LC-MS) in data-independent acquisition mode ^75,76^, we monitored the IOC2 expression in exponential and stationary growth phases (Methods M12), growing yeast in triplicates to compare the WT and mutant variant (n = 3 per group). We observed that the quantified IOC2 peptides of the mutant variant were on average ∼50% more highly expressed (Figure 3G) between S and E phases in comparison to the WT control (Methods 12), demonstrating that the mutant version of IOC2 extended the expression into the stationary phase in contrast to the wild type.

Finally, we analyzed the metabolic cost implications of abundance-increasing mutants compared to wild types, given concerns that increased protein copies might affect fitness ^19^. Overall, abundance-increasing mutant metabolic costs decreased significantly compared to random controls (Figure 3H, paired t-test, p-value < 1e-16). The most notable reductions were in synthesis under fermentative growth (*wagner_ferm*, −14% average) ^54^ and biosynthetic steps from central metabolism to the resulting amino acid (*craig_steps*, −13% average) ^55^. Both factors had a strong inverse relationship with BERT attention (Figure 1B & Table S1) confirming that the embedded space ordering (Figure 2A) and the model’s attention indirectly pick up the same evolutionary phenomenon. The exceptions were the impact of the relative change of the amino acid requirement on the minimal intake of ammonium ^51^ (*yeast_nit_rel*, 11% increase on average), which had the lowest correlation with attention, and the impact of relative change of the amino acid requirement on the minimal intake of glucose ^51^ (*yeast_car_rel*, 2% increase on average, see Table S7 for a full list). In summary, the significant cost reduction observed is especially striking since neither the BERT model nor the MGEM procedure were specifically trained with cost as a factor. This suggests that the neural network inherently recognized the connection between sequence cost and protein abundance, aligning with earlier observations on the cost-effective metabolism of highly abundant proteomes^32^.

## Discussion

Intracellular protein levels are determined by a delicate interplay of synthesis, regulation, and degradation. Despite the vast codon variability seen both within and between species at the DNA level ^77,78^, the conservation of protein ortholog abundances across diverse evolutionary lineages suggests an evolutionary imprint on amino acid sequences ^16–18^. While intricate cellular dynamics play a role in immediate protein concentrations, it is likely that significant evolutionary information resides within the primary sequence itself. Supporting this notion, the analysis of a consolidated proteomics dataset from a comprehensive list of yeast studies ^35^ showed that, while individual protein expressions vary, they mostly fluctuate around a specific value for 95% of proteins, but with the difference between proteins spanning over five orders of magnitude (Figure S1). This led us to postulate that amino acid sequences may inherently encode protein abundance. To explore this, we trained a deep neural network to predict protein abundance accounting for over half of the variability in abundance of the entire proteome dynamic range (Figure 1A, R^2^_test_ = 56%). By observing that amino acid composition across deciles of the dynamic range of protein expression is rather uniform (Figure S1), we confirmed that it is the amino acid arrangement in the sequence and not merely amino acid composition that is coding for protein abundance (Figure 1A inset).

The contributions of the various protein features on abundance have been studied mostly in isolation using linear models ^10,11,79^. However, given the dynamic nature of protein synthesis and degradation processes and their interactions, nonlinear models that integrate or abstract over the multiple levels are desired, especially given the loose coupling between some of these (e.g. the dynamic range of protein abundance is larger than that of mRNA and the former have longer half-lives ^79^). Thus, to decipher the biological insights gained by the neural network in predicting protein abundance, we analyzed the patterns within the BERT self-attention mechanism. Notably, attention profiles showed correlations with known protein abundance determinants (Figure 1B), including amino acid synthesis costs, suggesting that the model recognised the cell’s energetic currency concerning amino acid synthesis. The attention mechanism identified multiple associations between residues throughout the sequence, hinting at the neural network’s ability to discern overarching structural and physicochemical sequence patterns (Figure 1C). Our analysis further revealed that the network prioritizes regions with distinct secondary structure elements and functional domains when predicting protein abundance (Figure 1D, E). Moreover, the correlations found between attention, sequence structure, and physicochemical properties like polarity and hydrophobicity underscore the potential relationship between protein abundance and stability (Figure 1C).

The attention values in our model highlight crucial residue pairs for predicting protein abundance. While this theoretically points to specific sequence positions which are important for abundance prediction, understanding the encoder embedded space – a reflection of the sequence grammar grasped by BERT – is more challenging. This high-dimensional space encapsulates intricate sequence semantics and isn’t straightforward to interpret, resulting in a "semantic gap" between features and (human) meaning, often seen in deep learning models ^80,81^. To enhance our model’s explainability, we introduced the MGEM analytical framework. It simplifies the sequence space exploration by first establishing a one-dimensional reference (Figure 2A, B), then guiding mutations towards target sequence regions. Unlike methods that can produce unreliable predictions (predictor pathologies) ^82–84^ or local minima problems ^85^, MGEM deterministically modifies sequences based on their mapped target value, offering a deterministic solution for amino acid substitutions, beneficial for multiple applications. Furthermore, we believe this type of approach towards transparency and explainability of deep models warrants further work. As a future improvement, the procedure could be made free of guide sequences (and free of any bias towards these or inherent limitations stemming from the choice of the guide set), by constructing or training an inverse embedded-space-to-sequence mapping.

We applied the MGEM framework to perform a series of control-perturbation experiments to identify amino acids and protein properties that are intrinsically related to abundance (Figure 2A, B). In comparison to the random control that resulted in a decrease in protein abundance, MGEM-guided mutations achieved an average abundance prediction increase of over six times compared to the wild type sequences (Figure 2D). By inspecting MGEM mutants, we discovered that in terms of sequence position, the N-terminus was the most important, with the majority of amino acids remaining unchanged in this region (Figure 2E,F). This suggested that the N-terminus is generally evolutionarily optimized for expression efficiency, which also supports why it is widely used for protein expression optimization ^86–88^. A short hotspot at the very last position in the C-terminus was frequently mutated, which is known as a signal involved in protein degradation ^5,6^. Besides the C-terminus, however, most of the amino acids were substituted uniformly across the entire sequence length, mainly with the hydrophobic amino acids A (alanine), G (glycine) and V (valine) (Figure 2G). The introduction of hydrophobic amino acid residues into protein secondary structural components, such as helices, sheets and turns, is known to affect a protein’s conformational stability ^58,63,65^. We therefore hypothesized that there is a link between increased abundance and protein structure, and hence its stability.

We tested our hypothesis using extensive molecular dynamics (MD) simulations, an established technique for studying protein dynamics at the atomic level ^68,89^. Our data, derived from 200 MD simulations of random yeast proteins, showed that the majority of abundance-increasing mutations had increased the number of protein contacts and reduced solvent accessibility as reflected in reduced root mean square fluctuations (Figure 3A,D,E), phenotypes representative of stable proteins ^90–92^ (Figure 3D,E, Figure S6). The *in vivo* yeast proteomics experiment showed that these mutations resulted in sustained higher expression during growth phases (Figure 3G), further supporting our hypothesis that mutations increasing abundance also enhance protein stability. Note that here we kept codon frequencies the same as in the wild type strain, focusing solely on amino acid substitutions without modifying native gene regulatory regions, e.g. promoters. This approach likely leaves gene synthesis, transcription, and translation unaffected, while by observing long-term expression during the stationary phase, we assessed whether *in vivo* protein levels differed from the wild type due to changes in stability. While it is still unclear if the introduced mutations directly reduce *in vivo* protein degradation via stabilization of its conformation or operate through other mechanisms, our sequence perturbation experiments align well with previous observations that highly abundant proteins are generally more stable ^19,30,67,93^. This phenomenon is often explained by the so-called misfolding avoidance hypothesis and related hypotheses, which have dominated evolutionary discussions for the past decade, all aimed at explaining the slower evolutionary rates observed with highly abundant proteomes ^14,15^. An alternative explanation for the slow evolution of abundant proteins suggests that higher benefits come with higher costs ^15,33,34^. However, our findings indicate that proteins with mutations enhancing their stability are not only more abundant but also more cost-effective to produce. This explains their evolutionary advantage, as a structurally stable protein incurs fewer synthesis-associated costs to maintain consistent protein levels.

In conclusion, while the primary goal of our study was to investigate the relationship between a protein’s amino acid sequence and its abundance by examining a BERT network’s self-attention mechanism, our analysis revealed intricate connections between amino acid sequence, protein abundance, and metabolic cost related to protein stability. Remarkably, even without explicit conditioning on synthesis cost, both our BERT model and MGEM procedure succeeded in uncovering these latent relationships. This demonstrates the power of deep neural networks to decode complex biological systems. By manipulating the deep model’s semantics of these latent relationships, we unintentionally produced sequences optimized for cost. We demonstrate that mutations leading to increased abundance also contribute to enhanced protein stability, which in turn offers an evolutionary advantage by reducing the metabolic costs of protein synthesis. In addition, the MGEM approach opens new avenues in protein engineering by providing a robust, targeted method for amino acid substitution mapped to any continuous (real-valued) property. This has the potential for the design of proteins that are not only functionally efficient but also metabolically cost-effective, thereby offering a critical advantage in biotechnological applications. While no single theory can likely fully explain the complex relationships between protein sequence, abundance, and stability, our work identifies a critical link among these factors. By integrating insights from neural network predictions, extensive MD simulations, and *in vivo* experiments, we present a unified hypothesis that reaffirms the evolutionary advantage of stable, abundant proteins: they offer functional efficacy at a reduced metabolic cost.

## Methods

### M1. Neural Network Training

*Saccharomyces cerevisiae* (strain S288C) protein sequences were obtained from the UniProt^94^ reference proteome UP000002311 on 20th January 2020. To avoid technical challenges when training neural networks, we restricted the set of proteins to those with a length between 100 and 1000 residues (yielding 5202 out of 6049 proteins). The intersection of this set with the proteins with available abundance values from Ho et al. ^35^ resulted in 4750 unique sequences in our initial sequence-abundance dataset. To assemble the final dataset we added repeated measurements for each protein sequence, namely, each sequence appeared up to 21 times, each time with a different experimental target value from the Ho et al. dataset^35^, as in a regression with replicates, resulting in 99,603 training examples used as input/independent variable. Subsequently, for each sequence, a shuffled version was introduced with an “effective null” target value, a very small fractional value of 1e-5 (the unit for absolute abundance is molecules per cell), to allow for power transformations, resulting finally in 199,206 sequences. This was performed in order to expose the neural network to nonsense counter-example sequences so that it may learn to distinguish and to facilitate sequence interpretation, similar to training for classification problems ^95,96^ (here, with real and nonsense classes) or similar to using decoy sequences for distinguishing signal from noise in mass spectrometry 97. The data was randomly partitioned as 80% training, 10% validation, and 10% test, by splitting on unique sequences, i.e. ensuring repeated measurements of the same sequence were placed in the same data partition to avoid data leakage. Protein sequences (X’s / independent variable) and their corresponding target raw abundances (Y’s / dependent variable) were loaded as-is to BERT as input lists. To make the abundance distribution mass-centered, the preprocessing was configured to Box-Cox transform the raw abundances with λ = − 0.05155 using the expectation-maximization procedure as implemented in SciPy, on data based on medians of the initial dataset.

The training task’s preprocessing routine tokenized the sequences with the TAPE IUPAC^39^ tokenizer, each amino acid being assigned a unique integer value and the sequence flanked with special start and stop integer tokens. The TAPE^39^ implementation of the BERT *ProteinBertForValuePrediction* class was adapted for the model training. The model was trained as a regression task to minimize mean squared error (MSE). The model performance reported here was calculated by taking the median abundance across experiments for the proteins in the hold-out test set (436 values), as the test set obtained as above contained sequence repeats. The coefficient of determination was calculated on median values of the hold-out test using the Scikit-learn function. Hyperparameters search was performed using the BOHB algorithm ^98^ of the HyperBand scheduler ^99^ provided by the Ray library ^100^. Details about model architecture and hyperparameters are provided in Tables S9-S10. The best hypermodel thus found was then retrained. The best model consisted of 8 attention layers with 4 heads each (see Tables S8). The model was trained on a multi-GPU cluster using a mixture of A100 and V100 NVIDIA GPUs.

### M2. Attention profile analysis

As it is generally unclear ^101^ at which depth one might find lower or higher level features in such architectures, we considered all non-redundant attention profiles for a given sequence when measuring matches. Specifically, as BERT networks are known to have relatively high redundancy (i.e. different layers and attention heads learn very similar weights), we performed pairwise Pearson correlation of attention matrices from all layers and heads and kept only those that were uncorrelated (r < 0.01) with the majority (at least 90%) of other matrices, for each sequence. This left on average 4 non-redundant attention matrices per sequence. Moreover, attention matrices exhibited strong asymmetry (see Figure S2), often consisting of effectively uniform vertical streaks (i.e. the majority of residues “attend to” a single residue near-uniformly), thus making the “attended-by” values more informative (i.e. which residues receive such attention from all others). These “attended-by” values were averaged to produce one-dimensional attention profiles, which could be correlated with various per-residue measures. To match against qualitative data such as protein domains, we extracted residue attention *patterns* by keeping only the sequence positions that had an attention value z-score of at least 1 in the corresponding profile, to keep only those positions with the most signal.

### M3. Cost analysis

Per-residue cost profiles were computed for all proteins in the dataset (N = 4750) using the *S. cerevisiae* amino acid costs from Barton et al.^51^, with the exception of *yeast_sul_abs*, and *yeast_sul_rel*, which were deemed trivial for this task since they featured zero cost for all but a few amino acids. These profiles were then Pearson-correlated to all attention profiles for each protein (on average 4 attention profiles per protein), keeping only the maximum correlation with p-value < 1e-5 for each protein. The p-value was set using the Bonferroni correction for multiple testing at a target threshold of 0.05, thus resulting in 0.05 / 4750 = 1.053e-05.

### M4. AAindex Correlations

All 544 AAindex measures (https://www.genome.jp/aaindex, release 9.1 2006) were computed on a subsample of 1000 *S. cerevisiae* proteins using the R package Bio3D 2.4-3^102^. An average absolute correlation matrix was computed across the protein sequence subset and the AA indices were filtered using the R *findCorrelation* function (with a cutoff of 0.5) from the *caret* package 6.0-88, to only keep an non-redundant subset of 18 AA indices: BUNA790103, FINA910104, GEOR030103, GEOR030104, LEVM760103, MITS020101, NADH010107, NAKH920107, PALJ810107, QIAN880138, RICJ880104, RICJ880117, ROBB760107, TANS770102, TANS770108, VASM830101, WERD780103, WOEC730101. These per-sequence profiles for these indices were then computed for all proteins in the dataset (N = 4750) and Pearson-correlated to all attention profiles. Only the maximum correlation with p-value < 1e-5 was kept for each protein. The p-value was set using the Bonferroni correction for multiple testing at a target threshold of 0.05, thus resulting in 0.05 / 4750 = 1.053e-05. Note that the polar requirement (WOEC730101) was not part of the non-redundant list and was added manually due to its frequent description in the literature and the low correlation (r < 0.4) to the other indices. The resulting correlation distributions were filtered to only those AA indices with an absolute mean correlation of above 0.3 across all proteins.

### M5. Secondary structure analysis (DSSP)

Available *S. cerevisiae* PDB files (4745) generated by AlphaFold2 were downloaded from RCSB-PDB (on 2022-03-18). For each of these, DSSP 3.0.0 annotations were obtained using the BioPython 1.79^103^ *dssp_dict_from_pdb_file function*. For each protein and all its attention profiles (4 / protein, on average), DSSP annotations at positions with attention z-scores > 1 were counted. To avoid small numbers for significance testing, only structures with counts > 10 were kept. For all attention profiles, one-sided hypergeometric tests with a threshold p-value of 0.05 were performed both for enrichment and depletion of structure annotation counts, against the total background count of annotations across all proteins. Finally, this was summarized as the number of proteins that have attention profiles enriched or depleted in each type of DSSP structural annotation.

### M6. Domain analysis

Each InterPro domain was overlapped with the attention patterns produced for its protein (i.e. the positions of the sequence with attention z-score > 1), recording the highest overlap fraction (i.e. the largest fraction of *attended-to* domain residues) among all patterns produced for the sequence (output from all network layers and heads). To have a balanced control set, only domains that stretched to at most 50% of their protein length were kept (18,000 domains), so that the attention coverage inside the domain could be weighted against that outside of it. This was done (for each domain) by taking the number of high-attention positions outside the domain and dividing it by the number of times the domain could fit in the outside region (i.e. the number of windows the same length as the domain). This yielded an expected count corresponding to repeatedly randomly sampling subsequences the same length as the domain. The coverage fractions were taken as the the number of high-attention positions (either in the domain or the expected value outside) divided by the length of the domain. To assess the significance of the difference in domain coverage fraction distribution between attention and control, we performed a two-sided Wilcoxon signed-rank test, separately for each domain member database. The adjusted p-values were < 0.05 for 10 out of 12 member databases, where SFLD and HAMAP differences were not significant.

### M7. GO term enrichment analysis

The GO enrichment analysis for domains that overlap with attention was performed considering the proteins that have well-covered domains (>= 30% of their positions overlapping attention patterns) against the full set of proteins, with the Python library GOATOOLS 1.0.15^104^ using the Holm-Bonferroni p-value correction method and a significance threshold of 0.05. To summarize the results, GOATOOLS was used to obtain yeast GO slim terms (Table S4).

### M8. Embedded Ordering

To assess how individual amino acids in a sequence affect the abundance prediction, we probed the embedded space that the BERT encoder maps to. We call an *embedded ordering* the parametric UMAP projection ^105^ that we trained to map from this space down to a one-dimensional scale. The encoder’s embedded space contains 1024-dimensional point clouds (one cloud for each sequence) (Figure 2A), with every amino acid being assigned a (1024-dimensional) point. And because BERT uses a learned positional encoding, each residue in the sequence may be assigned a different value depending on position (i.e. regardless of the type of amino acid). From this space, a relatively simple feed-forward network (2 weight-normalized linear layers) is used for predicting values on the real line (Box-Cox-transformed protein abundances). The fundamental assumption of our construction is that (good) training induces a structure on the embedded encoder space that reflects the total order of abundance values (i.e. all scalar values are comparable and arranged in a strict succession). Under this assumption, we posit there exists a relatively low-dimensional manifold on which a geodesic connects all points in the (full) embedded space, resulting in an arrangement from lowest-prediction-value point clouds to highest-prediction-value point clouds (Figure 2A). The geodesic thus gives a total order within the embedded space. To retrieve a manageable approximation of the geodesic (and thus, of the order), we trained a parametric UMAP projection down to one-dimensional space. The embedded ordering thus constructed assigns a scalar value to each residue in the sequence, reflecting its contribution to the prediction. Moreover, these scalar values reflect a global ranking across the entire sequence space, i.e. lower abundance sequences will have residues with overall low order values, and the converse for higher abundance sequences. This enables easy assessment of the importance of each residue and enables mutation procedures.

The training set for the parametric UMAP consisted of the embedded start token point of each sequence, as information from the entire sequence is “routed” through these network nodes in the attention layers, and 10% of these were kept as a hold-out test set. The training was performed over multiple values of the UMAP number of neighbors hyperparameter, spanning an inclusive range from 1% to 25% of the number of sequences in the training set (aiming to balance local versus global structure). The performance was evaluated as the Spearman correlation between the centroids of the UMAP-projected point clouds and the corresponding abundance targets over test sequences.

### M9. Mutation Guided by an Embedded Manifold (MGEM)

The guided mutation was performed by sorting the residues according to their embedded ordering value and selecting the lowest of these for substitution, a different number for each scheme: the lowest 2, 5, 10, and 20 residues in each sequence, as well as the lowest 10%, 20%, and 30% of residues in each sequence. The 10 highest abundance sequences were selected as guides. This gives a pool of 4480 points distributed on the higher range of ordering values, available for substitution. For each residue selected to be substituted, its order value was increased by a large value, set as the width of the interval containing 99% of the embedded ordering (UMAP-projected) values, intuitively inducing a large shift in contribution to the prediction. To obtain a substitute residue that would match this shifted value, the guide sequences were used. The residue with the closest ordering value to this shifted value in each guide sequence was then chosen as a substitution candidate. This substitution was repeated for 10 guide sequences, and the one resulting in the highest prediction increase was finally selected. Both for the guided and the random substitution, the leading M residue was avoided. Random control was performed by choosing random residues (the same number as for each respective scheme) and substituting them with random amino acids.

### M10. Molecular dynamics (MD) simulations

We randomly subsampled 100 proteins with an increased abundance of at least 100% (from the 20% mutation regime, Figure 2D), ignoring transmembrane proteins. We applied molecular dynamics (MD) simulations to 100 mutated non-membrane yeast proteins showing higher abundance (Figure 2D, 20% mutation regime). Structures were generated both for mutated sequences and their corresponding wild types using AlphaFold2^48^. The structures were generated utilizing the full big fantastic database (BFD) and all five CASP 14 models ^48^. For each sequence, the structures with the highest average pLDDT score were then selected for molecular dynamics simulations. Simulations were carried out using the GROMACS simulation package 2022 ^106–108^, the AMBER99*-ILDN force field ^109^ and the TIP3P water model^110^. The protein was centered in a dodecahedron box with 1 nm distance to the box’s boundaries, solvated and neutralized by adding ions. The energy of the solvated system was minimized using a steepest descent algorithm (steps = 50,000, emtol = 1000 kJ/mol/nm, emstep = 0.01). Afterwards, the system was equilibrated for 100 ps in an NVT ensemble followed by a 100 ps equilibration in an NpT ensemble. For the productive run an NpT ensemble was chosen using the Parrinello-Rahman barostat (ref_p = 1 bar, tau_p = 2 fs, compressibility = 4.5e-5 bar^(−1))^111^. The temperature was set to 300 K using the v-rescale thermostat (tau = 0.1)^112^. For all steps periodic boundary conditions were applied in all dimensions. For the simulations a leap-frog integrator^113^ with a time-step of 2 fs was chosen. Covalent bonds involving hydrogens were constrained using the LINCS algorithm (lincs_iter = 1, lines_order = 4)^114^. Short range non-bonding interactions were cut off at 1 nm. For the van-der-Waals interactions a Verlet-cutoff scheme (ns_type = grid, nstlist = 10 steps, DispCorr = EnerPres), for the electrostatic interactions a Particle-Mesh-Ewald summation (pme_order = 4, fourierspacing = 0.16 nm)^115^ was applied. For each mutant and WT version of proteins, simulations were run for 100 ns. Protein coordinates were written to file every 1 ps. Simulations were considered converged if the RMSD was within a 10% error margin for 80% of the time points in the final quarter (Figure S8). Only these converged simulations (entire 100 ns) were selected for RMSF profile comparisons (Figure 3A).

### M11. Analysis of MD simulations

For the analysis, first, the periodic boundary conditions were fixed, and afterwards, the frames were rotationally and translationally fitted onto the protein atoms of the last frame of the trajectory using a least-square fit as implemented in GROMACS *gmx trjconv*. RMSF values were extracted using the GROMACS simulation package. Solvent accessible surface area (SASA) was computed using the implementation in GROMACS gmx sasa. The fraction of native contacts (Q2) were calculated from the last frame of the trajectory using the Python module MDAnalysis 2.2.0 ^116,117^. Contacts were defined as pairs of residues with an alpha carbon distance of 8Å or less. For the calculation of the DSSP^60^ and the solvent accessible surface area^118^ for the analysis of the protein UniprotID:Q12072 python package *MDTraj* 1.9.7 ^119^ was used. Dynamics were analyzed using VMD 1.9.4 and ChimeraX 1.4 ^120–122^. The structural images shown in Figure 3 were made with VMD. VMD is developed with NIH support by the Theoretical and Computational Biophysics group at the Beckman Institute, University of Illinois at Urbana-Champaign.

### M12. Proteomics analysis

The *S. cerevisiae* IOC2 knockout strain (*ioc2*Δ::*kanMX*) in the BY4741 (MATa *his3*Δ1 *leu2*Δ0 *met15*Δ0 *ura3*Δ0) background was requested from the Yeast Knockout (YKO) Collection ^123^ in Gothenburg University and used for genomic engineering in the following procedures. Predicted mutant (UniprotID: Q12072) DNA sequences flanking with 90 bp overlap to the specific genome sites on both ends were ordered as gene fragments from either TWIST Bioscience (www.twistbioscience.com). The mutant DNA sequence was designed such that it does not change original wild type codons to minimally affect the translation. The predicted mutated amino acids were substituted using most frequent corresponding codon.

To replace the *kanMX* gene ^123^ with the mutant gene in the genome, a gRNA plasmid targeting *kanMX* was constructed based on an All-In-One plasmid pML104 ^124^. The 20 bp gRNA sequence targeting at the *kanMX* gene (GCCGCGATTAAATTCCAACA) was designed with the CRISPR tool in Benchling (https://benchling.com). Primer sets pFA6-KanMX 488-507 FWD / pML_F and pFA6-KanMX 488-507 REV / f1 ori_R (Table S11) were used to amplify pML104 into 2 fragments pML104.part1 and pML104.part2 with 20 bp homologous sequences on both ends and gRNA sequence integrated in the pFA6-KanMX 488-507 FWD / pFA6-KanMX 488-507 REV primers. pML104.part1 and pML104.part2 were ligated into a circular plasmid named as pML104.gRNA_kanMX by Gibson Assembly ^125^ and was sequence-verified by Eurofins (https://www.eurofins.com/) with M13R primer (Table S11). pML104.gRNA_kanMX and mutant gene was transformed into knockout strain with PEG/LiAc method ^126^ and selected on synthetic minimal medium without uracil (SD-URA) plates. Colonies were verified with PCR using the primer set YLR095C_F / YLR095C_R (Table S1), and the amplified fragments were sequence-verified by Eurofins (https://www.eurofins.com/) with YLR095C_F / YLR095C_R primer set. SD medium supplemented with 5-fluoroorotic acid (SD+5-FOA) ^127^ was used to select colonies for loss of pML104.gRNA_kanMX.

Recombinant colonies without plasmids and the wild type BY4741 colony were picked into YPD medium. After overnight growth, 1% was inoculated into 1.5 ml YPD medium in a 48 well flower plate (M2P labs) and each sample had triplicates. The 48 well flower plates were cultured in 30 ℃, 1200 rpm for either around 10 h in a Biolector (M2P labs), until the cell growth reached mid-exponential phase, or 24 h until the cell growth reached stationary phase. 1 ml cells from both phases were collected and washed with MilliQ water once. After centrifugation, the supernatant was removed and cell pellets were kept in −80 °C until send to perform proteomics analysis at High Throughput Mass Spectrometry Core Facility, Charité (Berlin, Germany). Data independent acquisition was performed using the TimsTOF PRO mass spectrometer (Bruker) was coupled to the UltiMate 3000 RSL (Thermo). The peptides were separated using the Waters ACQUITY UPLC HSST3 1.8 µm column at 40°C using a linear gradient ramping from 2% B to 40% B in 30 minutes (Buffer A: 0.1% FA; Buffer B: ACN/0.1% FA) at a flow rate of 5 μl/min. The column was washed by an increase in 1 min to 80% and kept by 6 min. In the following 0.6 min the composition of B buffer was changed to 2% and column was equilibrated for 3 min. For MS calibration of ion mobility dimension, three ions of Agilent ESI-Low Tuning Mix ions were selected (m/z [Th], 1/*K*0 [Th]: 622.0289, 0.9848; 922.0097, 1.1895; 1221.9906, 1.3820). The dia-PASEF windows scheme was ranging in dimension m/z from 400 to 1200 and in dimension 1/K 0 0.6– 1.43, with 32 x 25 Th windows with Ramp Time 100 ms. Data quantification was performed using the DIA-NN 1.8 software, using library-free mode. Q12072 protein’s expression analysis in exponential and stationary phases (Figure 3G) was carried out using only the peptides that were detected in both growth phases in mutant and wild types correspondingly, i.e. the protein changes are calculated as fold-changes of corresponding Q12072 measured peptides in each strain. For the expression experiment three biological replicates from mutant and wild type were analyzed (6 samples in total). The raw mass spectrometry data have been deposited to the ProteomeXchange Consortium via the PRIDE partner repository ^128^ with the dataset identifier PRIDE:XXXXXXX.

### M13. Statistical analyses

All statistical analyses were performed using the Python (3.9) package Scipy 1.8.1^129^ and R 4.2.0. For data manipulation and visualization we used pandas 1.4.0 ^130^, seaborn 0.12.2 ^131^, scikit-learn 0.24.2 ^132^, and the R tidyverse 2.0.0 ^133^ package collection. Hypothesis testing was performed using the non-parametric Wilcoxon Rank Sum test, unless indicated otherwise.

### M14. Data and Software Availability

Scripts, training parameters, and software versions are provided in the following repository: https://github.com/fburic/protein-mgem

The models and data required to reproduce figures are stored in the following Zenodo record: https://doi.org/10.5281/zenodo.8377127

## Acknowledgements

The study was supported by SciLifeLab fellows program (A.Z.), Swedish Research council (Vetenskapsrådet) starting grant no. 2019-05356 (A.Z.), Formas early-career research grant 2019-01403 (A.Z.), WALP Wallenberg Launchpad project 2021.0198 supported by the Knut and Alice Wallenberg Foundation (A.Z.). A.Z is a Marius Jakulis Jason foundation scholar. The computations and data handling were enabled by resources provided by the National Academic Infrastructure for Supercomputing in Sweden (NAISS) and the Swedish National Infrastructure for Computing (SNIC) at the Chalmers Center for Computational Science and Engineering (C3SE), the National Supercomputer Centre in Sweden (NSC) and at the High Performance Computing Center North, partially funded by the Swedish Research Council through grant agreements no. 2022-06725 and no. 2018-05973. We thank Mikael Öhman and Thomas Svedberg at C3SE for their technical assistance. We thank Peter Dahl in Gothenburg University for offering the YKO strain. We thank Dr. Xiang Jiao in Chalmers University of Technology for providing the pML104 plasmid. We thank Dr. Gyorgy Abrusan and Dr. Oriol Gracia Carmona for discussions and feedback on the manuscript.

