## Supplementary Materials for "The amino acid sequence determines protein abundance through its conformational stability and reduced synthesis cost"

*These authors contributed equally

### Contents

Supplementary Figures 2
Supplementary Tables 9
Supplementary References 19

### Supplementary Figures

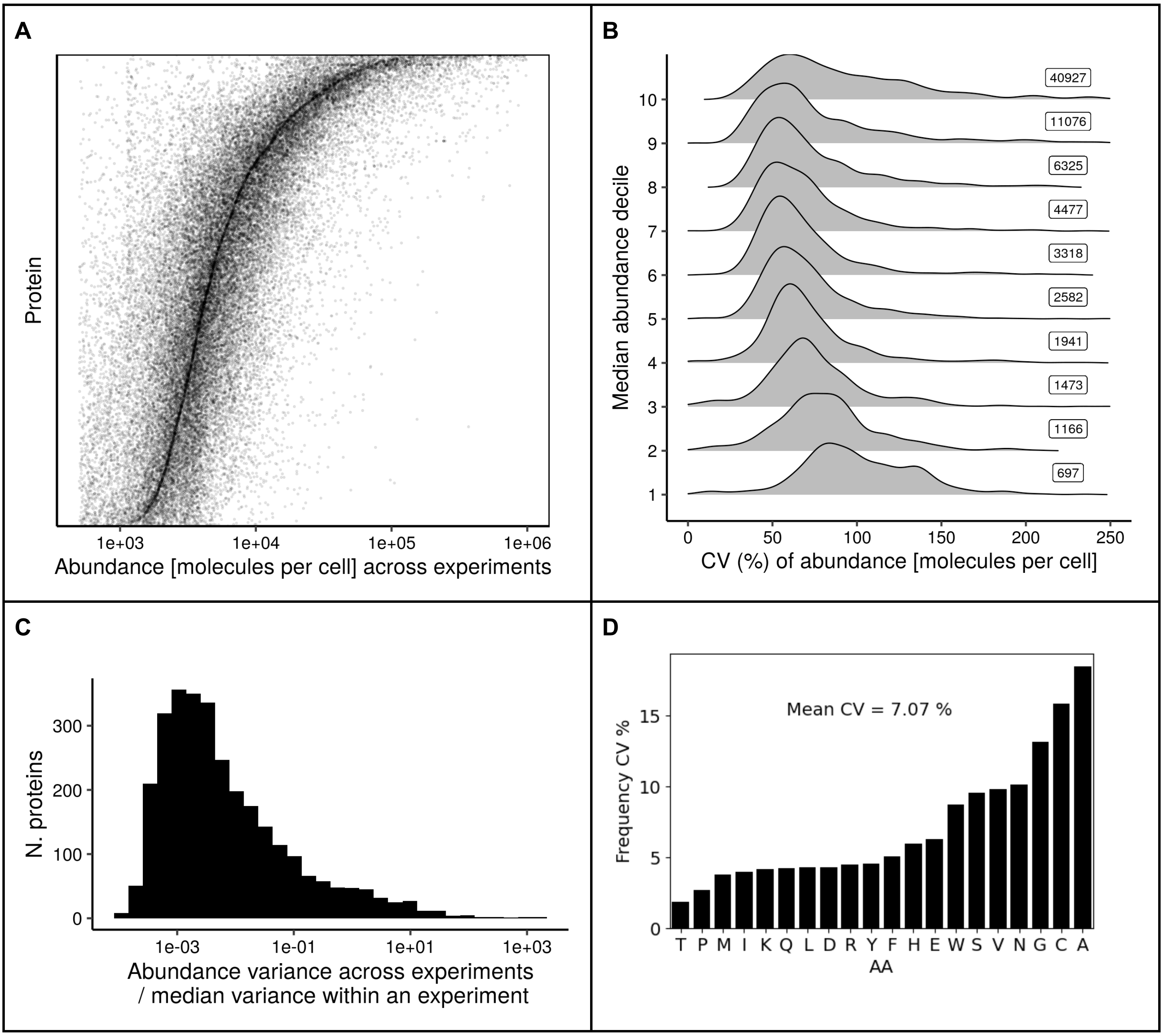


**Figure S1. Protein abundance and amino acid frequency have constrained variance.**
**A)** Protein abundance variation from data in (Ho, Baryshnikova, and Brown 2018)), showing the narrow variation across experiments for each protein (proteins with fewer than 10 experiment values were discarded)**. B)** Distribution of the coefficient of variation of protein abundance across experiments, grouped by abundance decile (right labels indicate median abundance for each decile). **C)** Distribution of ratios between the variance of protein abundance across experiments and the median of variances within each experiment. **D)** Coefficient of variation of amino acid frequencies across abundance deciles.

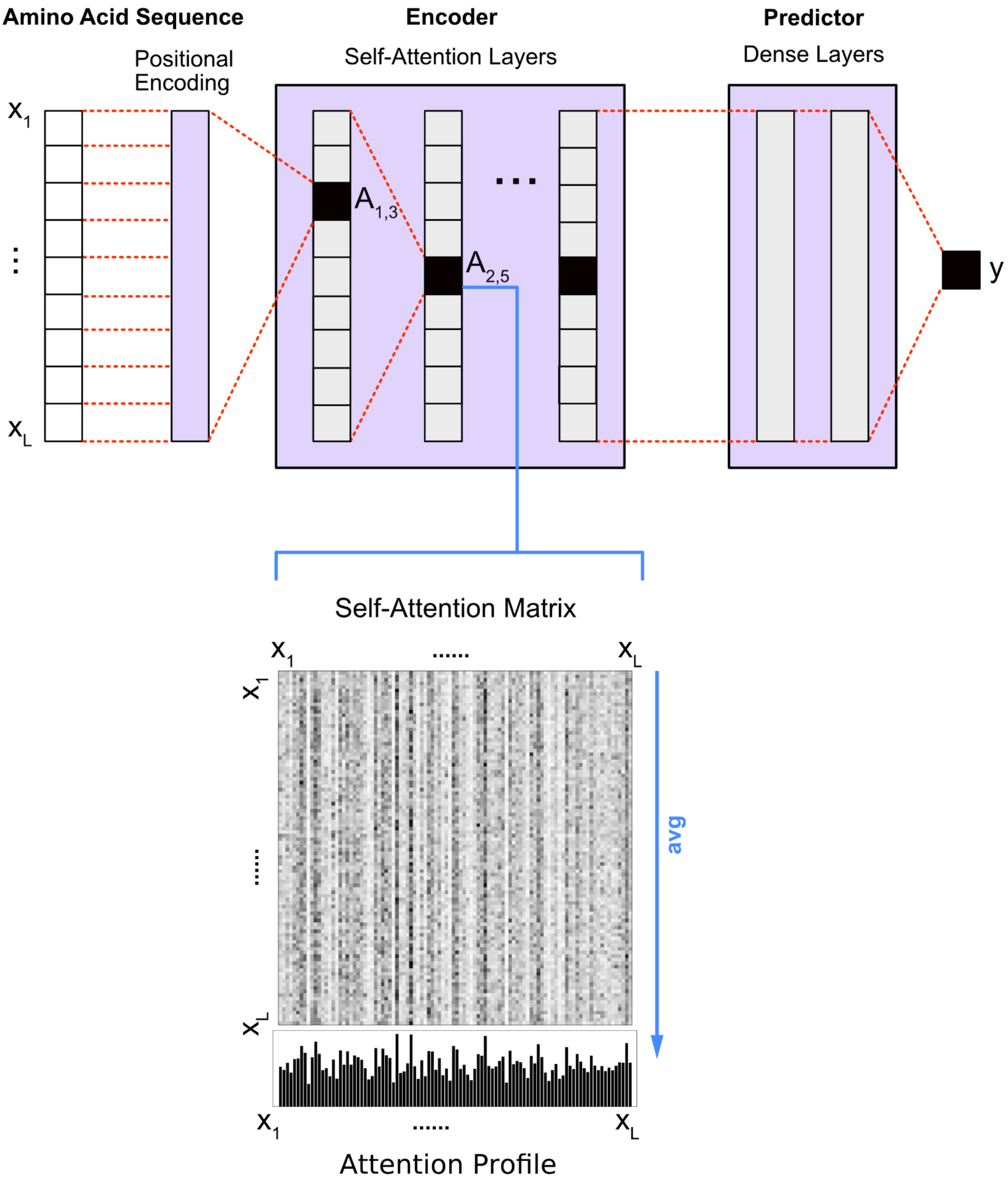


**Figure S2. Diagram of our BERT neural network (top) and examples of a self-attention matrix and derived attention profile (bottom).** Each head $A_{i,j}$ in an attention layer i outputs an attention matrix consisting of directional association weights between pairs of residues in the amino acid sequence (X, of length L), normalized as a percentage (across the entire matrix). The one-dimensional attention profiles used in this study were obtained by averaging along the "attends-to" axis, as the "attended-by" variation is generally more informative. A high value in a profile position signifies that the position "receives a lot of attention" from all other positions. Note that for a given sequence, one obtains N x H such matrices, for N attention layers and H heads.

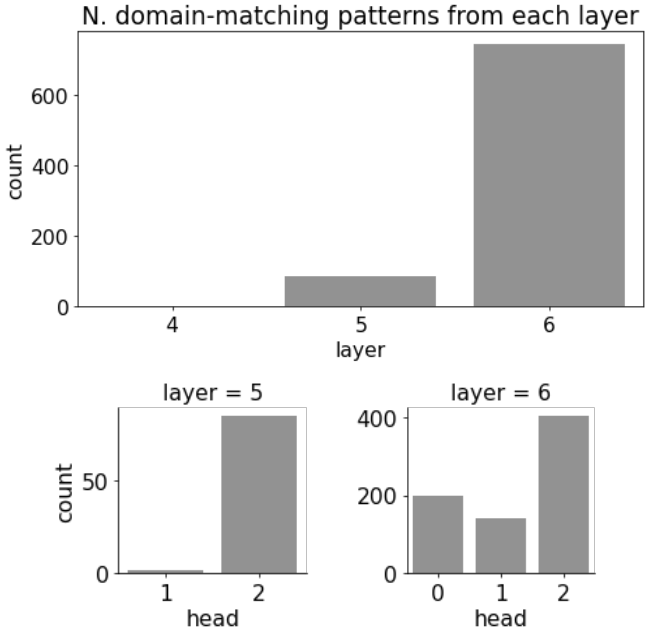


**Figure S3. Number of attention patterns matching protein domains, by layer and head.** The counts show the number of attention patterns that matched a protein domain in more than 30% of its residues. These patterns were obtained almost exclusively from the attention matrices of only a few heads, in the deeper attention layers of the BERT network (layers are numbered from 0 to 7).

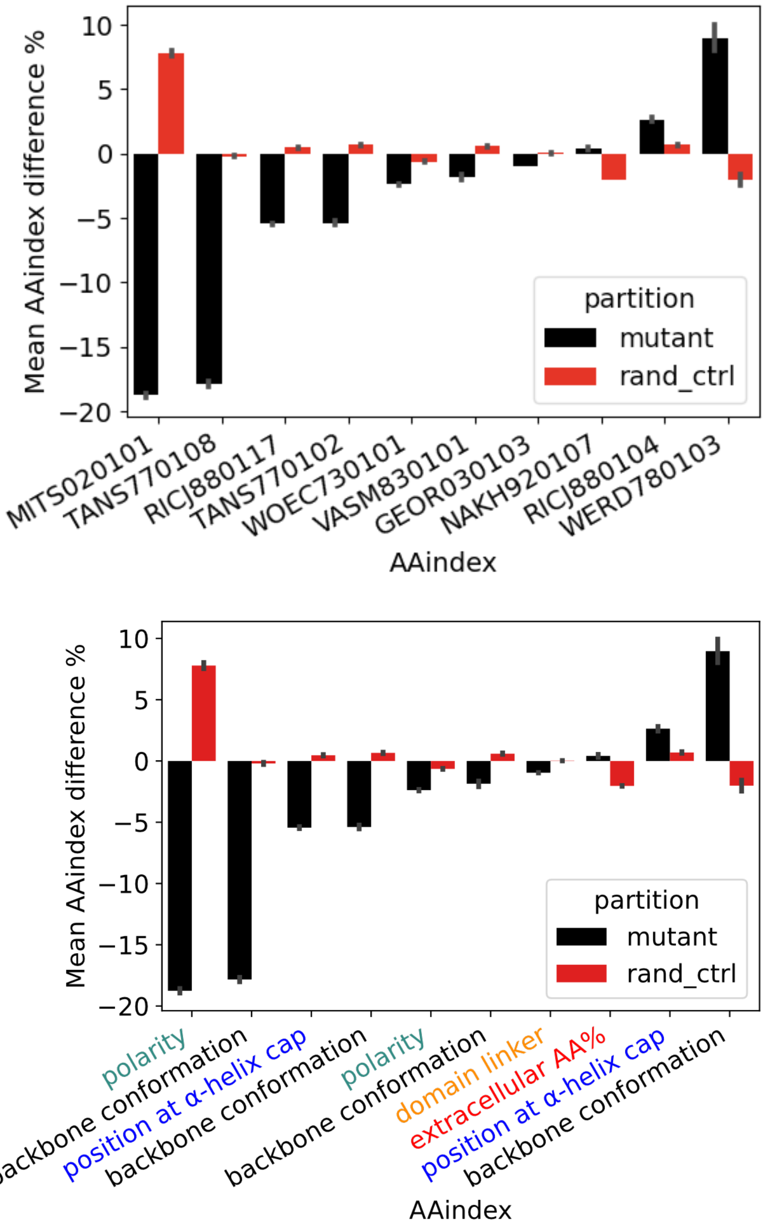


**Figure S4. The average sequence AAindex value of mutants obtained with MGEM show significant shifts compared with random control (paired t-test, p-value < 1e-308). Top**: AA indices are labeled by their AAindex IDs. **Bottom**: The indices are labeled by their type.

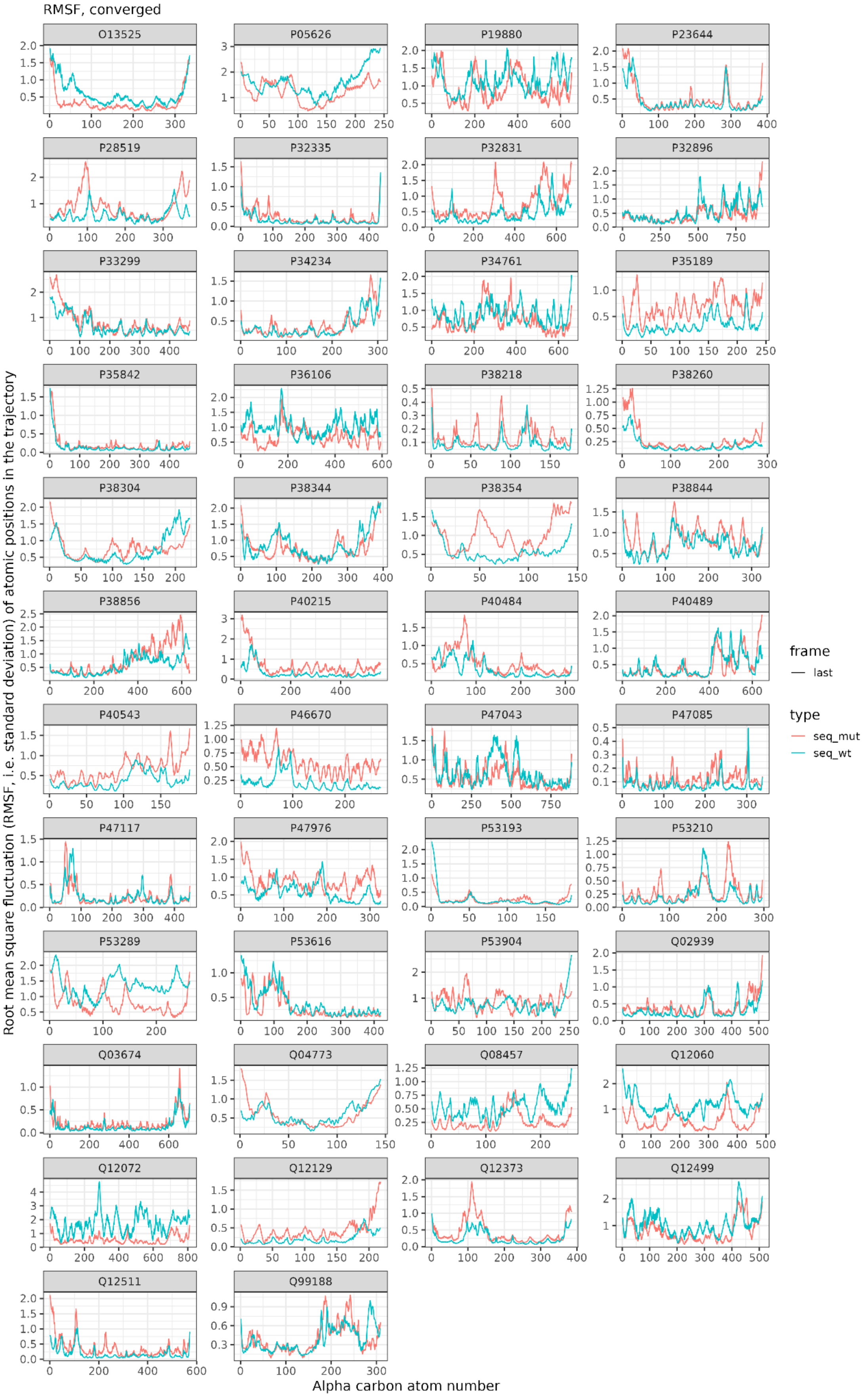


**Figure S5**. Root mean square fluctuations over 100 ns of MD simulations referenced to the last frame. Shown only converged simulations.

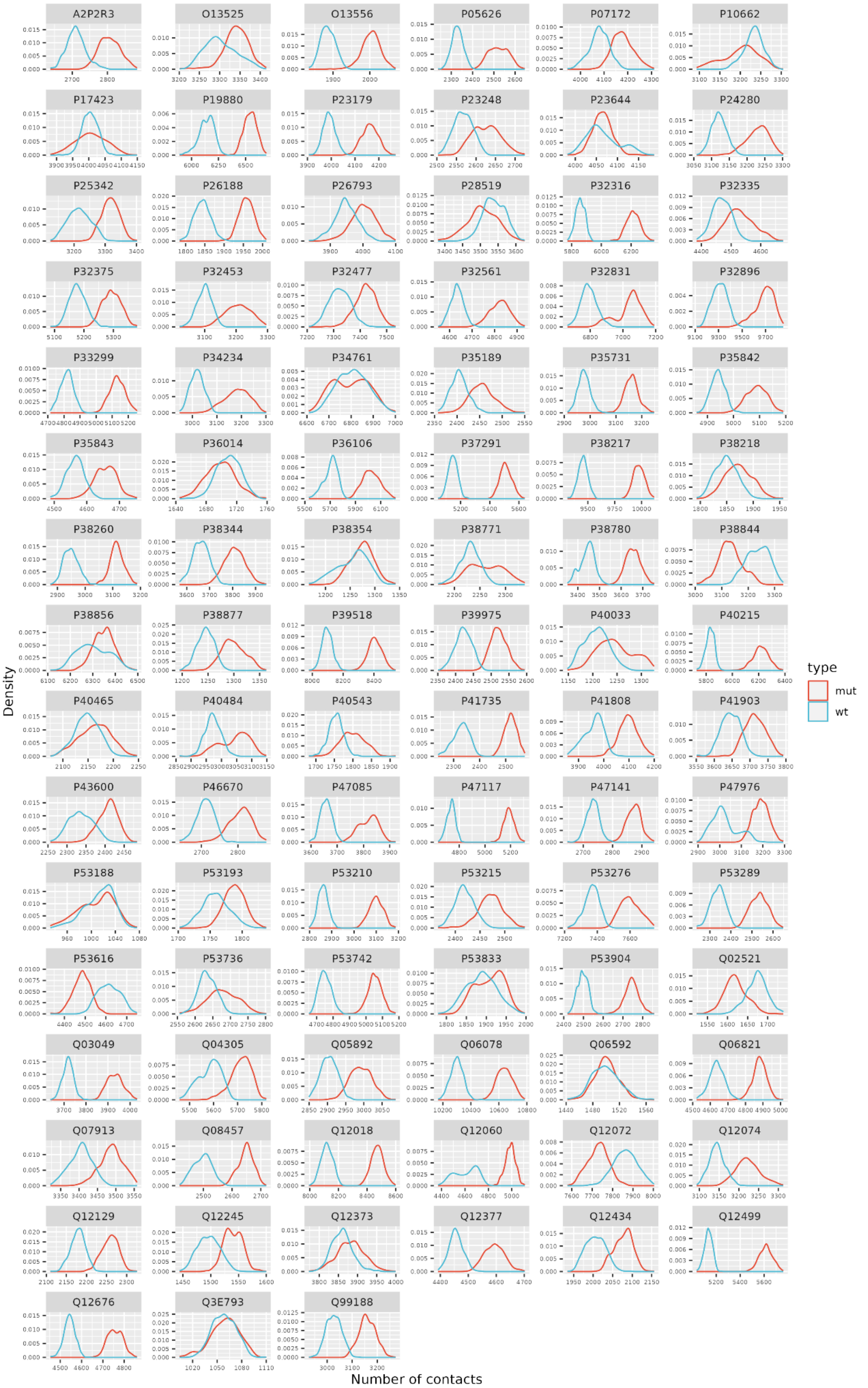


**Figure S6. Contacts Number contacts over simulation trajectory.** The contact is defined within 8Å distance between alpha carbons (see Methods).

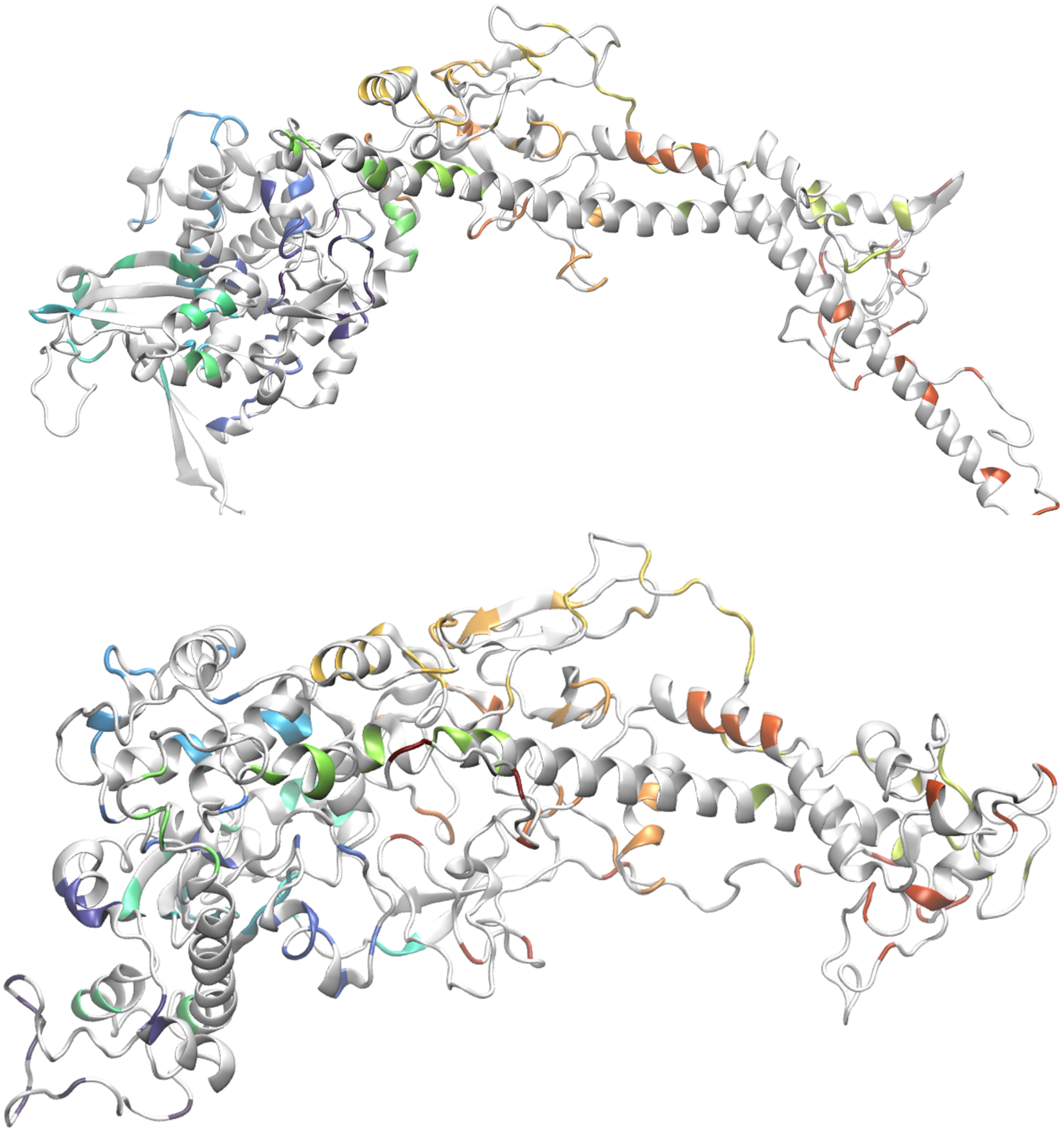

**Figure S7.** Mutated positions are highlighted in colours. The top panel denotes the last frame of the simulation of the wild type, the bottom panel of the mutatnt. The coloring is according to the amino acid index as shown in Figure 3F in the main text.

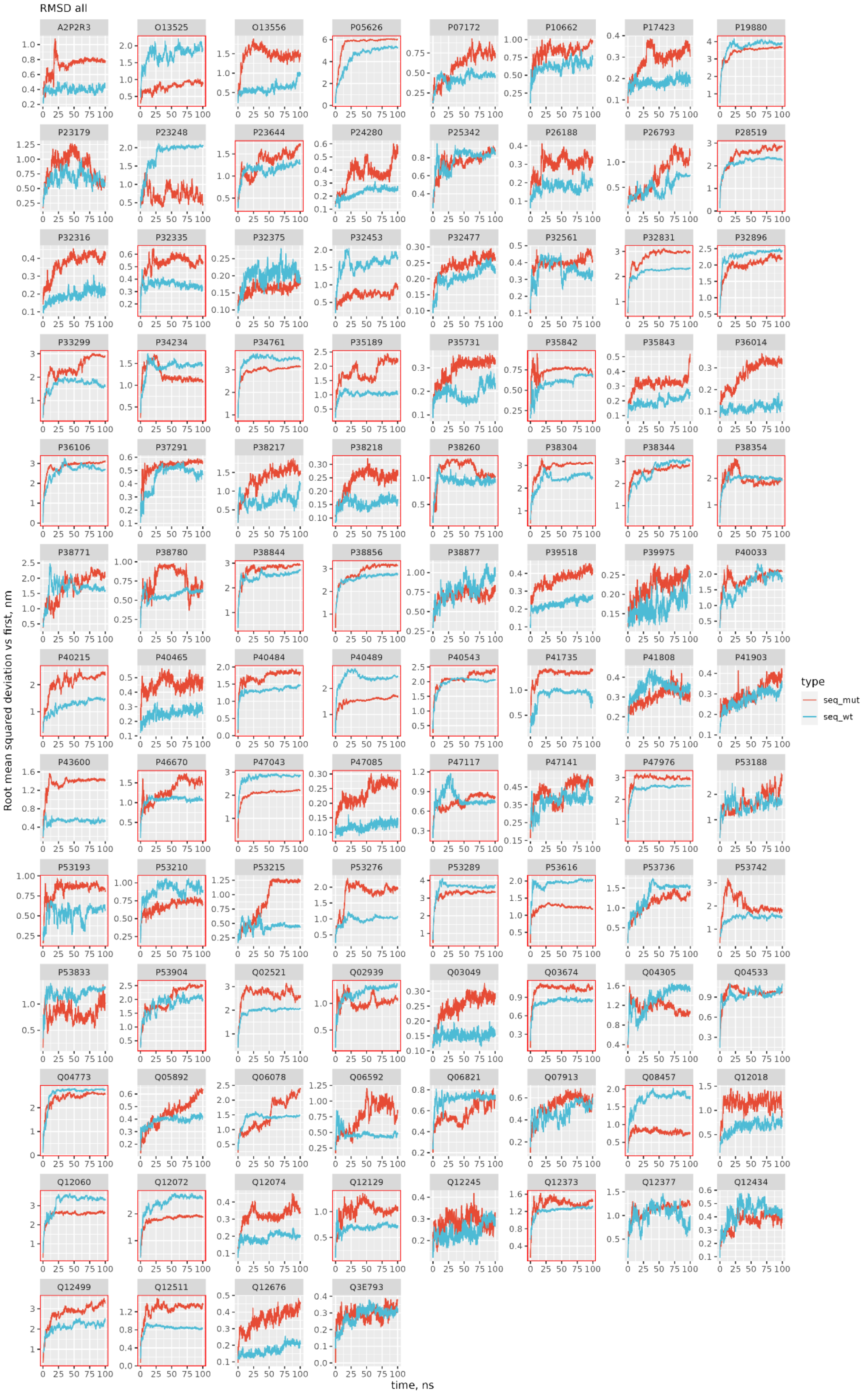


**Figure S8** Root mean squared deviation from first frame over 100ns of MD simulations. Highlighted are converged simulations where last quarter is within 10% of deviations (see main text Methods).

### Supplementary Tables

**Table S1. Amino acid costs that correlate with attention profiles.** The maximum absolute Pearson correlation (with p-value < 1e-5) was chosen among all the attention profiles of a given sequence. The table separates the positively and negatively correlated sequence subpopulations, showing mean values and subpopulation counts.

| Cost | Mean pos. corr. | Pos. count | Mean neg. corr. | Neg. count | Description |
|---|---|---|---|---|---|
| yeast_car_rel | 0.322725 | 1855 | -0.328086 | 705 | Impact of rel. change of the AA requirement on the minimal intake of C (glucose) (Barton et al. 2010) |
| craig_energy | 0.319014 | 1848 | -0.209371 | 684 | Energetic cost (avg. n. units of high energy P bonds and reducing H atoms required to produce the AA from glucose) (Craig and Weber 1998) |
| wagner_resp | 0.309522 | 32 | -0.346841 | 4226 | Cost of synthesis under respiratory growth (Wagner 2005) |
| wagner_ferm | 0.309046 | 66 | -0.329704 | 4057 | Cost of synthesis under fermentative growth (Wagner 2005) |
| yeast_nit_abs | 0.248459 | 1833 | -0.272822 | 1165 | Impact of abs. change of the AA requirement on the minimal intake of N (ammonium) (Barton et al. 2010) |
| akashi | 0.244109 | 33 | -0.286920 | 3341 | Energetic cost (avg. n. units of high energy P bonds and reducing H atoms) (Akashi and Gojobori 2002) |
| yeast_car_abs | 0.241446 | 121 | -0.289392 | 2797 | Impact of abs. change of the AA requirement on the minimal intake of C (glucose) (Barton et al. 2010) |
| weight | 0.230809 | 1438 | -0.325723 | 1472 | Molecular weight of the amino acid (proxy for synthesis cost) (Seligmann 2003) |
| yeast_nit_rel | 0.230152 | 1422 | -0.289971 | 4 | Impact of rel. change of the AA requirement on the minimal intake of N (ammonium) (Barton et al. 2010) |
| craig_steps | N/A | 0 | -0.311740 | 3932 | The number of biosynthetic steps between central metabolism and the resulting AA (Craig and Weber 1998) |

**Table S2. AAindex variables that correlate with attention profiles.** The maximum absolute Pearson correlation (with p-value < 1e-5) was chosen among all the attention profiles of a given sequence, for each profile. Shown are the mean correlation values across all proteins. AAindex values were fetched from https://www.genome.jp/aaindex (release 9.1 2006).

| AAindex | Mean corr. | AAindex type | Publication |
|---|---|---|---|
| RICJ880104 | 0.368029 | preference for position at α-helix cap | (Richardson and Richardson 1988) |
| WOEC730101 | 0.348570 | polarity | (Woese 1973) |
| MITS020101 | 0.305923 | polarity | (Mitaku, Hirokawa, and Tsuji 2002) |
| TANS770108 | 0.304256 | backbone conformation propensity | (Tanaka and Scheraga 1977) |
| VASM830101 | 0.301944 | backbone conformation propensity | (Vasquez, Nemethy, and Scheraga 1983) |
| NAKH920107 | -0.119901 | extracellular AA% (measured in membrane proteins) | (Nakashima and Nishikawa 1992) |
| RICJ880117 | -0.168564 | preference for position at α-helix cap | (Richardson and Richardson 1988) |
| GEOR030103 | -0.266115 | domain linker propensity | (George and Heringa 2002) |
| TANS770102 | -0.337806 | backbone conformation propensity | (Tanaka and Scheraga 1977) |
| WERD780103 | -0.381838 | backbone conformation propensity | (Wertz and Scheraga 1978) |

**Table S3. AAindex variables that correlate with attention profiles, separated in positively and negatively correlated protein subpopulations.** The maximum absolute Pearson correlation (with p-value < 1e-5) was chosen among all the attention profiles of a given sequence, for each profile. Shown are the mean correlation values across all proteins. AAindex values were fetched from https://www.genome.jp/aaindex (release 9.1 2006).

| AAindex | Mean pos. corr. | Pos. count | Mean neg. corr. | Neg. count | AAindex type |
|---|---|---|---|---|---|
| TANS770108 | 0.383633 | 3863 | -0.337209 | 482 | backbone conformation propensity |
| WERD780103 | 0.376552 | 8 | -0.383365 | 4359 | backbone conformation propensity |
| VASM830101 | 0.369531 | 3874 | -0.373924 | 386 | backbone conformation propensity |
| RICJ880104 | 0.369205 | 4018 | -0.231578 | 7 | preference for position at α-helix cap |
| NAKH920107 | 0.352343 | 1348 | -0.328829 | 3029 | extracellular AA% (measured in membrane proteins) |
| WOEC730101 | 0.348830 | 4389 | -0.224522 | 1 | polarity |
| GEOR030103 | 0.319206 | 270 | -0.310555 | 3565 | domain linker propensity |
| MITS020101 | 0.310673 | 3740 | -0.245472 | 32 | polarity |
| RICJ880117 | 0.295534 | 1016 | -0.329225 | 2944 | preference for position at α-helix cap |

**Table S4. GO slim terms for *S. cerevisiae* proteins that have domains captured by the model attention mechanism.** The GO slim terms were mapped from the significant GO enrichment analysis terms (Holm-Bonferroni-corrected p-value < 0.05).

| Biological Process | Molecular Funcion | Cellular Component |
|---|---|---|
| generation of precursor metabolites and energy | unfolded protein binding | cytoplasm |
| nucleobase-containing small molecule metabolic process | transferase activity | cytoskeleton |
| tRNA aminoacylation for protein translation | lyase activity | mitochondrion |
| protein folding | ATP hydrolysis activity | endoplasmic reticulum |
| response to chemical | kinase activity | membrane |
| translational elongation | ATPase-dependent activity | |
| chromatin organization | translation factor activity, RNA binding | |
| protein targeting | cytoskeletal protein binding | |
| regulation of translation | GTPase activity | |
| sporulation | peptidase activity | |
| protein dephosphorylation | glycosyltransferase activity | |
| protein phosphorylation | oxidoreductase activity | |
| monocarboxylic acid metabolic process | transmembrane transporter activity | |
| carbohydrate metabolic process | hydrolase activity | |
| | DNA binding | |
| | phosphatase activity | |
| | ion binding | |
| | helicase activity | |
| | ligase activity | |

**Table S5. The leading 30% of protein sequences in the yeast proteome differs in composition from the overall sequence.** The amino acid counts of 4750 proteins were computed for the leading 30% region and for the entire sequence. These counts were compared using one-sided hypergeometric tests for enrichment and depletion, with a threshold p-value of 0.05.

| Enriched AA | p-value | Depleted AA | p-value |
|---|---|---|---|
| A | 9.679602e-04 | C | 3.849427e-15 |
| H | 2.059567e-03 | D | 3.104653e-08 |
| M | 5.330124e-76 | E | 1.043352e-49 |
| P | 7.746553e-22 | F | 1.895432e-15 |
| Q | 1.768194e-10 | G | 2.204858e-08 |
| R | 3.393047e-08 | I | 1.693833e-22 |
| S | 2.381427e-128 | K | 8.558290e-08 |
| T | 4.209202e-24 | L | 6.875738e-09 |
| | | V | 4.648197e-04 |
| | | W | 3.563835e-44 |
| | | Y | 2.146192e-09 |

**Table S6. Mean AAindex difference for optimized proteins obtained with MGEM.** The control consisted in (uniformly) randomly selecting replacement amino acids for the same number of positions as the corresponding MGEM mutant (avoiding the leading Met). AAindex values were fetched from https://www.genome.jp/aaindex (release 9.1 2006).

| AAindex | Mean mutant AAindex difference % | Mean random control AAindex difference % | Publication |
|---|---|---|---|
| MITS020101 | -18.753839 | 7.789475 | (Richardson and Richardson 1988) |
| TANS770108 | -17.832835 | -0.200313 | (Woese 1973) |
| RICJ880117 | -5.443304 | 0.464476 | (Mitaku, Hirokawa, and Tsuji 2002) |
| TANS770102 | -5.388635 | 0.659881 | (Tanaka and Scheraga 1977) |
| WOEC730101 | -2.380900 | -0.645872 | (Vasquez, Nemethy, and Scheraga 1983) |
| VASM830101 | -1.854750 | 0.595889 | (Nakashima and Nishikawa 1992) |
| GEOR030103 | -0.957061 | 0.023577 | (Richardson and Richardson 1988) |
| NAKH920107 | 0.413022 | -2.036270 | (George and Heringa 2002) |
| RICJ880104 | 2.637145 | 0.691204 | (Tanaka and Scheraga 1977) |
| WERD780103 | 8.963559 | -2.015681 | (Wertz and Scheraga 1978) |

**Table S7. Mean cost difference for optimized proteins obtained with MGEM.** The control consisted in (uniformly) randomly selecting replacement amino acids for the same number of positions as the corresponding MGEM mutant (avoiding the leading Met).

| cost | Mean mutant cost difference % | Mean random control cost difference % |
|---|---|---|
| wagner_ferm | -13.603179 | 3.933833 |
| craig_steps | -13.306743 | 1.682922 |
| akashi | -9.218666 | 3.059934 |
| yeast_car_abs | -8.899055 | 1.926197 |
| wagner_resp | -8.736643 | 2.193350 |
| weight | -6.159937 | 0.863105 |
| craig_energy | -5.551885 | 3.734348 |
| yeast_nit_abs | -2.709100 | 1.179117 |
| yeast_car_rel | 1.978379 | -2.667477 |
| yeast_nit_rel | 11.187093 | -2.451302 |

**Table S8. The architecture of the BEST model with best performance.** The implementation was the TAPE *ProteinBertForValuePrediction* class, consisting of positional encoding, an encoder made up of multi-headed attention layers, and instead of a decoder (as in typical Transformer architectures), a thin (2 dense layers) multi-layer perceptron (MLP) predictor to a real value. To process our protein data, we implemented a specific TAPE learning task class to be used with the model.

| BERT schematic | Network element | Value |
|---|---|---|
| 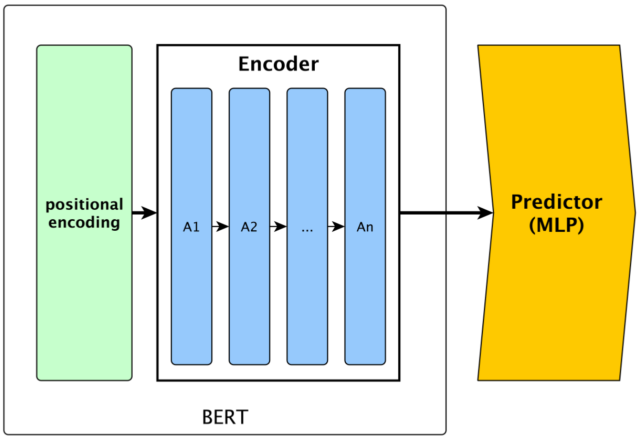 | vocab_size | 30 |
| | type_vocab_size | 2 |
| | max_position_embeddings | 1024 |
| | num_hidden_layers (attention layers) | 8 |
| | num_attention_heads | 4 |
| | hidden_size (encoder embedding dim.) | 1024 |
| | hidden_dropout_prob | 0.0 |
| | attention_probs_dropout_prob | 0.0 |
| | hidden_act | relu |
| | initializer_range | 0.02 |
| | layer_norm_eps | 1e-12 |
| | intermediate_size (MLP) | 3072 |



**Table S9. BERT Training parameter values.** These were used for retraining the best model found by the hyperparameter search.

| Training Parameter | Value |
|---|---|
| learning_rate | 2.5681e-07 |
| batch_size | 32 |
| gradient_accumulation_steps | 16 |
| num_train_epochs | 500 |
| patience | 50 |

**Table S10. Hyperparameter search space for BERT models.** The hyperparameter search was performed with Ray Tune using the *HyperBandForBOHB* scheduler (HyperBand that enables the BOHB Algorithm). Dropout was effectively disabled by fixing the hyperparameter range to zero. The Ray hyperparameter search used 10 samples per trial (choice of hyperparameter values).

| Hyperparameter | Search space |
|---|---|
| learning range | [1e-8, 1e-6] (uniform log-space sampling) |
| number of training epochs | {500, 800} |
| batch size | {16, 32, 64} |
| number of hidden layers | {8, 10, 12, 14} |
| number of attention heads | {4, 8, 16} |
| hidden size | {512, 1024} |
| intermediate size | {2048, 3072, 5120} |
| hidden dropout probability | 0 |
| attention dropout probability | 0 |
| hidden activation function | {gelu, relu} |

**Table S11.** List of PCR primers.

| Primer name | Sequence |
|---|---|
| pFA6-KanMX 488-507 FWD | GCAGTGAAAGATAAATGATCGCCGCGATTAAATTCCA ACAGTTTTAGAGCTAGAAATAGC |
| pFA6-KanMX 488-507 REV | GCTATTTCTAGCTCTAAAACTGTTGGAATTTAATCGC GGCGATCATTTATCTTTCACTGC |
| pML_F | ACGCGCCCTGTAGCGGCGCA |
| f1 ori_R | TGCGCCGCTACAGGGCGCGT |
| M13R | CAGGAAACAGCTATGACC |
| YLR095C_F | GACACGACTAAGAAACTAGATCAATTGCTC |
| YLR095C_R | TACGGATGTGCGGCTGGAAAAGAAAG |
